# StORF-Reporter: Finding Genes between Genes

**DOI:** 10.1101/2022.03.31.486628

**Authors:** Nicholas J. Dimonaco, Wayne Aubrey, Kim Kenobi, Amanda Clare, Christopher J. Creevey

## Abstract

Large regions of prokaryotic genomes are currently without any annotation, in part due to well-established limitations of annotation tools. For example, it is routine for annotation tools to misreport or completely omit genes using alternative start codons. Therefore, we present StORF-Reporter, a tool that takes an annotated genome and returns missing CDS genes from unannotated regions. StORF-Reporter consists of two parts. The first begins with the extraction of unannotated regions from an annotated genome. Next, Stop-ORFs (StORFs) are identified in these unannotated regions. StORFs are open reading frames that are delimited by stop codons and thus can capture those genes most often missing in genome annotations.

We show that this methodology recovers genes missing from canonical genome annotations. We inspected the results of the genomes of model organisms, the pangenome of Escherichia coli, and a further 6,223 prokaryotic genomes of 179 genera from the Ensembl Bacteria database. StORF-Reporter was able to extend the core, soft-core and accessory gene-collections, identify novel gene families and extend families into additional genera. The high levels of sequence conservation observed between genera suggest that many of these StORF sequences are likely to be functional genes that must now be added to the canonical annotations.

## INTRODUCTION

Prokaryotic genomes are most often observed with high levels of gene density, resulting in low proportions of unannotated DNA, especially compared to Eukaryotes (1, 2). Much of this unallocated DNA is intergenic, i.e. found between annotated functional elements such as genes, promoters or other functionally and structurally important regions. In eukaryotes, the relative proportion of a genome reported to be intergenic has been linked to the biological complexity of the organism (3). However, in prokaryotes intergenic regions are often assumed not to be functionally important, despite studies reporting putative coding and non-coding genes (complete and pseudogenised) within (4, 5, 6).

Recently, evidence has emerged of elevated levels of conservation and selection in putative intergenic regions (7) suggesting that these regions should be investigated to determine whether they contain genes or functional elements missed by contemporary prediction methods. Therefore, in this work we investigate the potential of prokaryotic ‘intergenic regions’, (referred from this point on as <u>unannotated regions</u> (URs) for clarity) to harbor yet-to-be discovered CDS genes.

Methods employed to study URs often rely on sequence similarity to known proteins deposited in databases and as such are limited to the set of proteins deposited there and by the cut-offs used to identify a significant match, especially when attempting to identify a gene fragment from an UR. Therefore, to elucidate these regions and overcome these and other limitations inherent in identifying coding regions in prokaryotic genomes (2), we have developed a novel methodology to identify and extract putative CDSs in the URs of existing prokaryotic genome annotations. This approach, named ‘StORF-Reporter’, is performed in two stages. First, URs are extracted from an annotated genome (see Figure 1), then we identify and filter from these extracted URs open reading frames (ORFs) that are delimited by two in frame stop codons (termed StORFs for Stop-ORFs).

**Figure 1.**
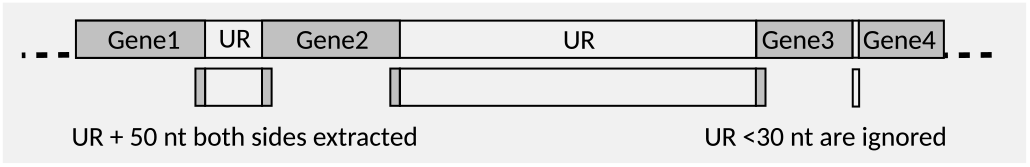
Visual representation of how unannotated regions (URs) are selected for extraction. URs that are less than 30 nt are not extracted. URs are extracted with an additional 50 nt on their 5’ and 3’ ends to allow for overlapping genes and the upstream untranslated region between the first stop codon and the true start codon.

Conflicting definitions of ORFs exist in the literature (8), and while it is the norm for prokaryotic annotation tools to annotate an ORF as the region between a start and stop codon in a coding sequence, it should be noted that the Sequence Ontology (9) makes no reference to a start codon in its definition of an ORF. Instead, it is described as “the inframe interval between the stop codons of a reading frame which when read as sequential triplets, has the potential of encoding a sequential string of amino acids.”. Our definition of StORFs is therefore synonymous with the definition of an ORF in the Sequence Ontology but for clarity and due to the prevalence of the start-to-stop codon definition of an ORF in the literature, we refer to stop-to-stop open reading frames as StORFs hereafter.

An example of a StORFcan be seen in Figure 2, which captures multiple possible start codons. The choice of which start codon to use as the start of the coding region in a situation like this will usually come down to the specific weightings inherent in the tools used and/or the matches they received to any databases that are used in the annotation process. This quite often results in non-standard start codons being underrepresented in prokaryotic annotations (2). Shifting the focus from locating the most likely start codon for an ORF to identifying the first upstream in-frame stop codon simplifies the problem as it reduces reliance on correctly identifying genomic features such as the Shine–Dalgarno motifs (10), promoter regions (11), ribosomal binding sites, and operons (12), and reduces the potential for inadvertently truncating the predicted protein product by choosing incorrectly between alternative putative start codons (see Figure 2A). This is important as while the canonical ‘ATG’ start codon is most-often used to initiate approximately 80% of prokaryotic CDS genes, many species and cross-species gene families have been shown to possess very different start codon profiles (13, 14). Furthermore, there is growing evidence of the existence of organisms that adjust their genetic code in response to changing environments (15), thus making traditional start codon identification even more complex. Unlike start codons, one of the three canonical stop codons (TGA, TAG, TAA) is the terminus for almost all CDS genes in almost all species, and despite the possible influence of purifying and/or positive selection (16, 17), they can be reliably used to identify the terminus of protein synthesis in a gene. Additionally, it has been observed that stop codons appear more frequently than expected in non-coding reading frames possibly due to selection against translation of the wrong frame following frameshift mutations or ribosomal slippage (18). Thus, the longest StORF identified from all 6 reading frames in any UR has the most potential to represent a coding region.

**Figure 2.**
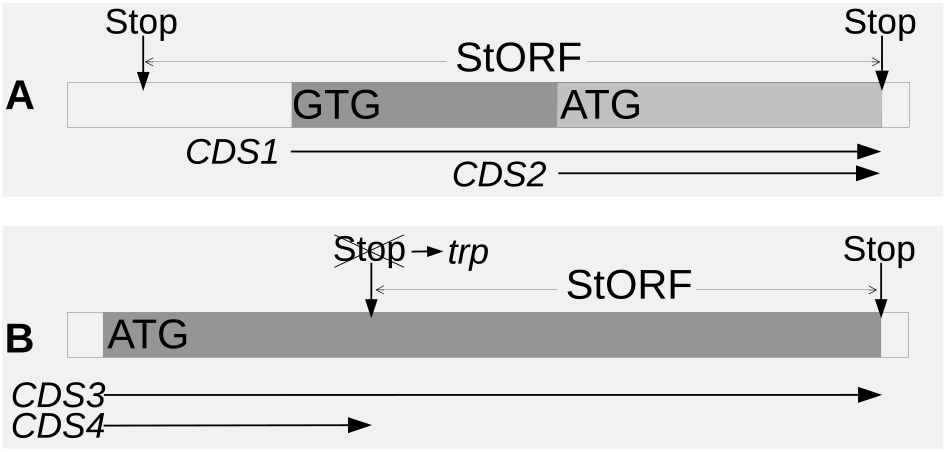
Visual representation of a StORF and how it can capture multiple potential start codons in an unannotated region. Part **A** depicts a StORF capturing two possible start positions (GTG and ATG) for a CDS gene which could produce two distinct CDS sequences (*CDS1* and *CDS2*). Part **B** shows how a StORF can comprise of only a partial segment of a gene if that gene either recodes a canonical stop codon (Stop to trp) or has had an in-frame stop codon mutation resulting in one complete and one truncated transcript (*CDS3* and *CDS4*).

We propose the StORF-Reporter methodology as an additional step to supplement contemporary prokaryotic genome annotations from *de novo* annotation workflows (such as PROKKA (19)) or from databases (such as Ensembl (20)) in order to identify overlooked putative coding regions and as such we provide tools to supplement existing genome annotation files with information on any StORFs identified. The resulting enhanced annotation builds upon the work that proceeded it rather than restarting annotations from scratch.

To demonstrate the utility of this approach, we identified and investigated StORFs from three increasingly complex sets of genomes. Firstly, we examined six bacterial model organisms which exhibit many URs long enough to contain full-length CDS genes, expanding the narrative of these well-studied genomes.

Secondly, we investigated the presence of conserved StORFs across the 219 *Escherichia coli* (*E. coli*) strains elucidating their impact on our understanding of the E. coli pangenome.

Lastly, we applied the methodology to 6,223 prokaryotic genomes representing 1,417 species from Ensembl Bacteria to understand the trends of StORFs across diverse bacterial groups.

## MATERIALS AND METHODS

### Datasets

The canonical annotation and sequence data for 44,048 genomes of 8,244 prokaryotic species, which included 43,347 bacterial genomes and 493 archaeal genomes, were acquired from the 46^th^ release of Ensembl Bacteria (20). For each genome, two data files were downloaded; the complete DNA sequence (*\*-dna.toplevel.fa*) and the GFF (Generic Feature Format) file (*\*.gff3*) containing the position information for each gene (coding and non-coding). The genomic elements (including both coding and non-coding genes) presented in the Ensembl GFF annotations were taken as reference annotations for this study. Fragmented genomes were removed by filtering out those with more than 5 contigs (not including plasmids). Then, genera that had less than 5 genomes were removed. This resulted in 6,223 genomes (see Supplementary Table 1). Out of this 6,223, the six model organisms; *Bacillus subtilis (B. subtilis*) BEST7003 strain, *Caulobacter crescentus (C. crescentus*) CB15 strain, *E. coli* K-12 ER3413 strain, *Mycoplasma genitalium (M. genitalium*) G37 strain, *Pseudomonas fluorescens (P. fluorescens*) UK4 strain, *Staphylococcus aureus (S. aureus*) 502A strain, shown in Table 1, were selected for the first study. Additionally, *E. coli* was selected to undergo a pangenomic analysis of its Ensembl annotated genes and extracted StORFs. As many of the Ensembl genomes were unlabeled at the species level, to make sure the correct genomes were extracted, only those labeled specifically as *‘Escherichia-coli’* were extracted. This resulted in a group of 219 *E. coli* genomes from the original filtered group of 6,223 (see Supplementary Table 2 for further detail).

**Table 1.** An overview of genome composition for the six model organisms used in the analysis of StORF-Reporter. This table shows the number of Ensembl-annotated genes (coding and non-coding). The gene density is presented in bold square brackets. These six organisms span a range of genome size (0.58 - 6.06 Mbp) and gene density (percentage covered with annotation, 83.93% - 92.03%).

| Model Organism | Genome Size (Mbp) | Genes [Density] |
| --- | --- | --- |
| *B. subtilis* (BEST7003) | 4.04 | 4,133 [88.91%] |
| *C. crescentus* (CB15) | 4.02 | 3,875 [90.60%] |
| *E. coli* K-12 (ER3413) | 4.56 | 4,257 [86.28%] |
| *M. genitalium* (G37) | 0.58 | 559 [92.03%] |
| *P. fluorescens* (UK4) | 6.06 | 5,266 [84.75%] |
| *S. aureus* (502A) | 2.76 | 2,556 [83.93%] |

### StORF-Reporter

StORF-Reporter consists of two distinct parts which can be used independently or together as they were in this study: URExtractor(.py) and StORF_Finder(.py). The processes of both are described in the following sections, and the code is available at https://github.com/NickJD/StORF-Reporter. A third tool, StORF_Reporter(.py) is available to perform UR and StORF extraction from PROKKA (19) annotated genomes and presents the user with a newly created PROKKA formatted GFF file for use in downstream tools that require the PROKKA format.

#### Unannotated Region - Extractor (UR-Extractor)

To facilitate the extraction of URs from different prokaryotic genomes (and the various interpretations of the GFF format), UR_Extractor was developed. Written in Python3 (21) and using user-defined genomic features in GFF format to identify the boundaries of genomic elements, regions of DNA without annotation were extracted from the provided DNA sequence file. The output consists of a GFF3 and FASTA file with the details of the sequence and loci for each UR that can be used further analysis (see Supplementary Section 2.1 for command line menu of UR_Extractor.py).

When recovering URs from a genome it is important to consider that StORFs may overlap with annotated genes (22). Therefore, it was necessary to determine the extent to which the URs should be extended into the annotated regions. The 6,223 genomes from Ensembl Bacteria, including the six model organisms in Table 1 were studied to identify representative parameters for the extraction of URs across this large set of Ensembl Bacteria genomes. The median CDS gene overlap observed for each of the six genomes was 3 nt and the 75^th^ percentile was below 50 nt, as can be seen in the left half (A) of Figure 3. The lengths of overlap between genes (both coding and non-coding) across the 6,223 prokaryotic genomes were investigated and, as can be seen in Supplementary Figure 1, the gene overlap lengths observed for the 6 genomes are similar to the majority of the Ensembl annotated genomes. Furthermore, the median distance from a gene’s canonical start codon to the nearest in-frame upstream stop codon (representing the untranslated region of a StORF) was 39 nt (see the right half (B) of Figure 3). These findings demonstrate that StORFs should contain very little extraneous upstream DNA.

**Figure 3.**
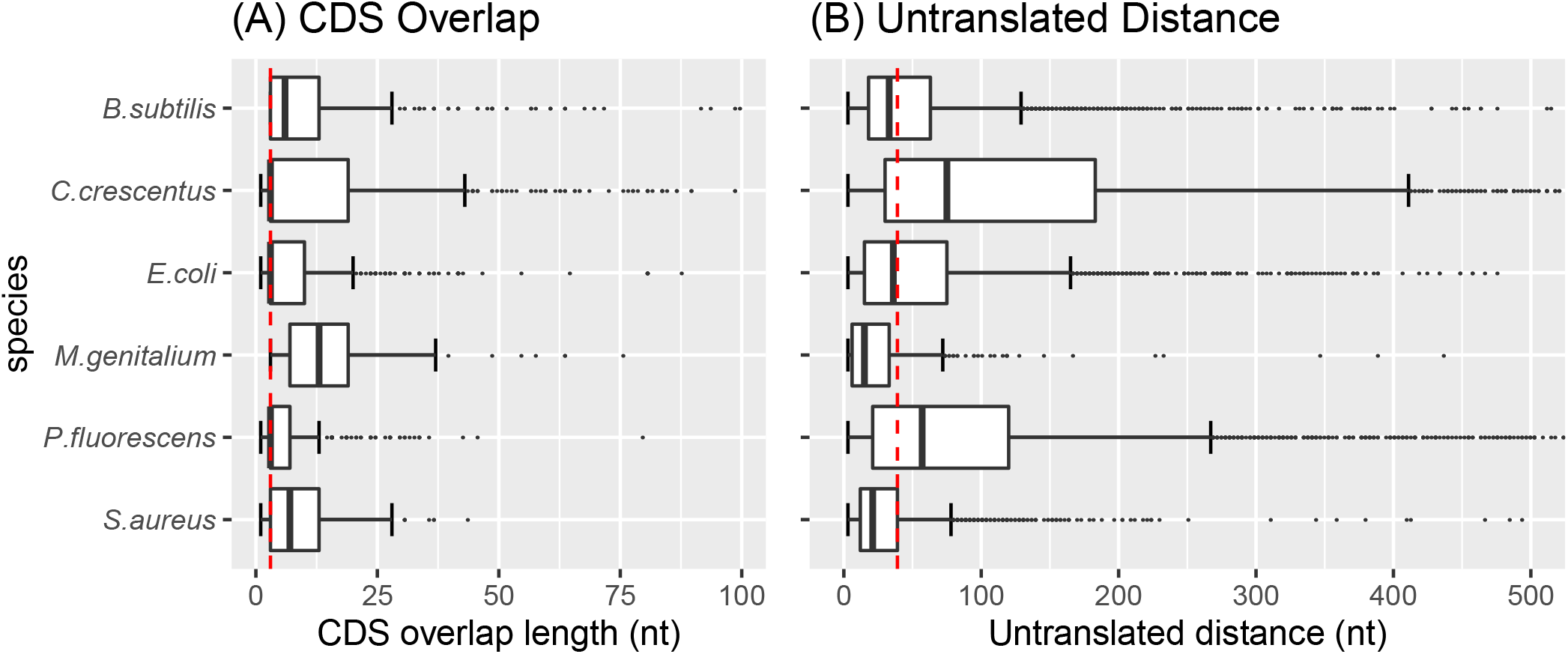
This double plot reports the analysis of the 6 model organisms which were used during the parameterisation of StORF-Reporter. Figure A reports the distributions of the Ensembl annotated CDS gene overlap lengths for each model organism with a dotted red line representing the overall median (3 nt) with the x-axis truncated at 100 nt. Figure B reports the distance between an Ensembl gene’s start codon and the first in-frame upstream stop codon for the model organisms with the x-axis truncated at 500 nt. The dotted red line represents the overall median (39 nt). These plots indicate that the extension of 50 nt from each end of the extracted intergenic regions is often enough to capture both the true overlap between an annotated gene and the putative gene identified by a StORF and the small amount of upstream non-coding DNA which the StORF will contain.

To account for the observed median gene and StORF features, URs were only extracted if they were at least 30 nt long but were extended by 50 nt at both the 5-prime and 3-prime ends (into annotated genes) by the UR Extractor tool (see Figure 1), totaling a minimum length of 130 nt. This extension of 50 nt at both ends was to allow for the capture of the parts of StORFs which may overlap with existing gene annotations. This process also took into account the untranslated region between the start of the StORF and putative start codon of the protein sequence it captured and was necessary as the direction of a potential StORF is not known in advance (see Figure 2 for an illustration of how a StORF captures upstream DNA).

Genomic repositories, including Ensembl Bacteria, are known to contain varying levels of assembly and annotation error due to a number of constraints (23, 24, 25). It can also be the case that genomes have missing annotations as a result of human or software error. Therefore, UR Extractor has a maximum UR length cutoff of 100 kb. A genome with a 100 kb UR either warrants further investigation for biological interest or is likely to contain annotation error.

The parameters used for UR extraction are available in Supplementary subsection 3.1.

#### Stop - Open Reading Frame - Finder (StORF-Finder)

To identify StORFs, StORF-Finder starts by scanning the previously extracted URs (or alternatively any DNA sequence) for loci encapsulated by in-frame, user-definable stop codons (see Figure 2). All three stop codons are frequently present in intergenic regions and also within genes but out-of-frame, hypothesised to prevent off-frame reading of gene sequences (26). To investigate whether StORF-Finder should preferentially select one stop codon over another, a comparison was performed on the abundance of the triplets with their usage as stop codons across the Ensembl genomes. The results of this analysis (see Supplementary Figure 2) show that while there are differences between the three stop codons, they are not universal and it is likely not beneficial to make generalisations.

As with start codons, stop codons are differentially preferred by groups of genes and gene-families both from within the same and across different species (27). For example, we can see in Supplementary Figure 2 that the codon TAA is often used more frequently than its distribution across the genomes suggests. Similarly, TGA is often used less frequently than its distribution across the genome would suggest in all 6 model organisms (with TGA never used as a stop codon in *M. genitalium* (28). For TAG, the usage is mostly in line with the overall distribution across the genomes. There is clear evidence of stop codon preference and even repurposing for amino acids in species such as *M. genitalium* and specific gene-types which utilise their T/UGA and T/UAG codons (29) for selenocysteine (30) and pyrrolysine (31). Nevertheless, these cases are rare and it can be assumed that for the majority of prokaryotic genomes, the three canonical stop codons can be treated equally.

While removing the need to identify the location of start codons, StORFs do have their own drawbacks. To be consistent with the universal genetic code, codon table 11, only the codon ATG is translated to methionine (M) in the StORF reporter output. This presents a problem when reporting StORFs since, alternative start codons (i.e. codons other than ATG which initiate translation) are incorrectly assigned the amino acid represented in the codon table 11 and not methionine as is observed in the protein (32). This is a difficult problem to overcome since we cannot predict alternative translation start codons in advance. To mitigate this, StORF-Finder can report both the DNA and amino acid sequences in its output.

As with traditional CDS prediction methods, without any filtering, StORF-Finder can report a very high number of overlapping and nested StORFs, even for short regions of DNA. Therefore, a filtering step was developed to reduce the number of StORFs reported. This is done by first ordering all StORFs by their length in descending order and then iteratively removing the nested or shorter StORFs in cases where two StORFs overlap beyond the defined length of 50 nt. This is done individually for each set of StORFs for each UR and is intended to maximise the likelihood that recovered StORFs contain functional sequences. The remaining StORFs are reported in FASTA (DNA and/or amino acid form) and GFF format with both locus coordinates for their original position within the original genome sequence, and their position relative to the UR in which they were reported.

The parameters used for StORF Finding are available in Supplementary subsection 3.2.

### Extracting StORFs from Prodigal and Ensembl annotations of the six Model Organisms

To investigate the ability of StORF-Reporter to add to and improve contemporary annotations, URs and StORFs were extracted from the 6 model organisms. This was done using the existing annotations provided by Ensembl Bacteria and for the annotations provided by running the Prodigal (33) genome annotation tool on all 6 organisms. The default parameters of URExtractor (-gene_ident “ID=gene”) were used to identify the complete set of genomic elements (coding and non-coding) which were then used to extract URs from the Ensembl provided annotations. Prodigal was applied to the complete DNA sequences of the six model organisms and a novel CDS prediction was performed using the tool’s default parameters. As Prodigal only predicted CDSs, UR_Extractor used the CDS coordinates to extract URs (-gene_ident “CDS”).

### *Escherichia coli* Pangenome Analysis

To investigate the impact that StORFs may have on the pangenome of a well-studied prokaryotic model organism, the Ensembl protein sequences and StORFs extracted from a collection of 219 *E. coli* strains were studied.

Although gene loci information is often used to build a pangenome, many of the URs which potentially harbor the novel CDS genes that StORF-Reporter identifies are unlikely to be in the same location across all genomes. Additionally, the fragmented state of the majority of genomes held in Ensembl Bacteria (as is the case for many repositories) further adds to the necessity to use alternative techniques to identify the pangenome. Amino acid sequence similarity is an established metric used to group gene families. It is used by the clustering tool CD-Hit, which does not require loci information (34). Amino acid sequences present in the *E. coli* genome were downloaded from Ensembl Bacteria (*.pep.all.fa*) and combined into one FASTA file and gene-clustering was then undertaken with CD-Hit (version 4.8.1) (34). Gene clustering was performed with the following parameters: aligned sequence identity threshold of 0.9 (90%), length difference cutoff of 0.6 (60%), and the ‘-g’ option was set to the ‘more accurate’ option (see Supplementary Subsection 3 for more details). The consistent use of the 0.6 length difference cutoff between clustered sequences allowed for the instances where the StORF sequences contained additional upstream non-coding DNA which would have hindered the clustering of the matching coding regions. The strict sequence identity threshold of 0.9 ensured that the resulting clusters were very similar across the regions where they did align.

The output of CD-Hit consists of two datafiles, a ‘.clstr’ (cluster) file containing the full sequence cluster metadata and a FASTA file containing the representative ID and sequence for each cluster. The CD-Hit cluster file, which consisted of the identified Ensembl gene clusters, was used as a baseline against which to compare the additional StORF sequences. The amino acid sequences of the *E. coli* StORFs recovered from the URs for the same set of 219 genomes were then combined with the previous CD-Hit output (FASTA) from the Ensembl proteins, which consisted of one representative sequence for each cluster (representatives are often the longer of the sequences in any one cluster). This combined FASTA file then underwent another round of CD-Hit clustering with the same parameters and produced the final *E. coli* pangenome datafiles (see Supplementary Figure 3 and Subsection 3.3 for more details on the files created and used in this process).

The results of these clusters are complex and have been classified differently with respect to the origins of each sequence set. As such they will be used throughout the rest of this paper and are referred to as follows:

- **Ensembl-Only**, which refers to the clusters with only Ensembl annotated protein sequences.
- **Ensembl-StORF**, which refers to the clusters which contain at least one Ensembl annotated protein sequence and one StORF amino acid sequence.
- **StORF-Combined-Ensembl**, which refers to the clusters where StORF sequences combined at least 2 or more Ensembl cluster representatives together.
- **StORF**, which refers to the clusters in the Ensembl-StORF group that were categories as core, soft-core or accessory with only the StORF sequences being counted.
- **StORF-Only**, which refers to the clusters which solely contain StORF amino acid sequences.

To interpret the output from CD-Hit, a number of Python3 scripts were developed and are available on the StORF-Reporter GitHub repository https://github.com/NickJD/StORF-Reporter. The tool **CD-Hit_StORFReporter_Pangenome_Builder.py** was built to first identify the Ensembl core-gene families for the 219 *E. coli* strains. Next, it then identifies whether any StORFs clustered with these representative gene cluster groups, but also whether there were any StORF-Only gene clusters spanning multiple *E. coli* strains. This allows for the recording of unique strains/genomes that make up each cluster and therefore the ability to calculate each cluster’s pangenomic status (core, soft-core, accessory). The StORF-Finder output sequence ID tag ‘Stop-ORF’ was used to distinguish between the Ensembl and StORF sequences.

It is difficult to predict what the result of clustering many thousands of protein sequences together could be. Through introducing hard-coded computation parameters into the natural world, it is inevitable that certain cases are unaccounted for. For example, multiple representatives from separate Ensembl gene clusters could be combined into a new single cluster with the addition of StORF sequences. To account for this, the Ensembl clusters that were combined when StORF sequences were added were first recorded separately in the Ensembl gene cluster results, and then additionally recorded in their new combined clusters with the additional StORFs (see results rows 1 and 2 respectively in Table 3).

The parameters used for the CD-Hit clustering are available in Supplementary subsection 3.3.

### Extracting StORFs from 6,223 Ensembl genomes

An important aspect of gene family research concerns their diversity and spread across different genera. The StORF-Reporter methodology was applied to the 179 genera collection of 6,223 filtered Ensembl Bacteria genomes using the Ensembl annotations for the UR and StORF extraction. The diversity of assembly and annotation quality of these genomes presented a number of obstacles. As with the *E. coli* genome collection, even after filtering, the genomes in this study were often in a fragmented state with a number of low quality regions containing large sections of ambiguous nucleotides. With this in mind, URs and StORFs were extracted individually for each of the 6,223 genomes using default parameters. The resulting StORF-Finder outputs were then combined into a single FASTA file with their original genus and species name appended to the start of each sequence header. For example, the full genome name of ‘Enterobacter_cloacae_ecwsu1.asm23997v1’ was appended to the start of the protein name ‘AEW71445’ to make ‘Enterobacter_cloacae_ecwsu1.asm23997v11AEW71445’ The same CD-Hit parameters and workflow were undertaken as for the *E. coli* pangenomic study. A modification of the previously developed single species script was built to handle multiple genera. This script records multiple levels of taxa information for each of the sequences such as genera, species and strain. However, as the analysis of the *E. coli* pangenome had already demonstrated StORF-Reporter’s ability to capture gene sequences at the species level, only the genera specific to each sequence are reported in this study.

### Validation of recovered StORFs

Different validation methods were required to determine what was being detected by the StORF sequences. For the StORFs identified from within the URs of the Prodigal annotations, an extension to the ‘ORForise’ platform (2) was developed, which originally was designed to compare annotations from different CDS prediction tools to a reference annotation. This extension, ‘StORForise’, was used to identify the Ensembl CDS genes missed by Prodigal that StORF-Reporter was able to recover. The default parameters of what classified a gene as ‘detected’ were taken from the ORForise package (a minimum of 75% coverage of an in-frame CDS prediction for a reference CDS gene). StORF-Reporter was only presented with the extracted URs according to the Prodigal CDS predictions. Therefore, mispredictions by Prodigal could result in elongation of an Ensembl CDS gene prediction or prediction of a CDS where no Ensembl annotation existed. These would constrain the available regions that could be searched for by StORF-Reporter.

Therefore, as part of the ORForise platform, two additional scripts were developed to identify the Ensembl genes that Prodigal missed and specifically the number of missed Ensembl genes which were non-vitiated (not corrupted or contaminated) by the mispredictions of Prodigal (false positives, overlapping Ensembl genes by more than 50 nt).

The protein sequence aligner Diamond (35) (version v0.9.30.131) was used to align the recovered StORFs from the Prodigal and Ensembl annotations of the six model organisms to known proteins from both the same genomes proteome and to the Swiss-Prot protein database (36) (release 2021 _01, downloaded 05/03/2021). The sequence similarity alignments were performed with the Diamond blastp option with two separate parameter runs, one with a minimum bitscore of 60 and the second with the same bitscore but also a subject coverage cut-off of >=80%. This was done to report whether some StORFs were as a result of gene duplication or whether they had previously been identified in other studies. This was performed at the amino acid level as although the coding start sites are not identified by StORF-Finder, the coding frame is, and so the upstream region will include the start codon in the same frame (unless mutations are present). This indicates that StORFs can be directly translated into amino acid sequences, and then undergo similarity analysis to find homologous proteins already reported across other genomes and protein databases.

## RESULTS

### Unannotated regions of model organisms

UR-Extractor was first applied to both the canonical Ensembl annotations and the Prodigal predictions of the six model organisms. Details of these URs can be seen in Supplementary Tables 3 and 4. These show that the numbers of URs reported for each model organism were similar between the Ensembl and Prodigal annotations, except for *M. genitalium* which were 157 and 636 respectively. This was likely due to the inability of Prodigal to account for the re-purposing of the ‘UGA/TGA’ stop codon causing a number of predicted CDSs to be incorrectly truncated. Additionally, the Ensembl annotations also included non-coding genes which were treated the same as coding genes by the UR-Extractor tool. As Prodigal does not predict non-coding genes, it is likely to report additional or longer URs than those recovered using the Ensembl annotations. This can explain some of the much longer UR lengths in the Prodigal analysis for the other model organisms. For example, while the longest UR length in *S. aureus* was 2,591 in the Ensembl annotation, this increased to 12,332 in the Prodigal annotation. However, this was not consistent across all model organisms. Interestingly, the longest UR in P *fluorescens* decreased from 20,088 to 4,264 for Ensembl and Prodigal annotations respectively.

There were clearly a number of URs that were long enough to contain complete CDS genes extracted from both the annotations of Prodigal and Ensembl - these were studied further.

### StORF-Reporter recovers Ensembl CDS genes missed by Prodigal

An analysis of the StORFs extracted from the URs of the six model organism genomes was undertaken using the annotations from Prodigal. StORF-Reporter was able to recover Ensembl-annotated genes that were missed by Prodigal, and also found many other potential genes that had sequence similarity to the Swiss-Prot database or proteins already annotated in the model organism genomes.

The ORForise platform reported that Prodigal was able to identify the vast majority of Ensembl genes from each of the model organisms, except for *M. genitalium* (2). For each of the model organisms, StORF-Reporter identified a high number of StORFs, as can be seen in Supplementary Table 5. However, StORF-Reporter is impeded by the mispredictions produced by Prodigal. For each model organism, Prodigal reported a number of CDSs which either had no equivalent in the Ensembl annotation, were elongated versions of Ensembl genes or were too inaccurate to be classified as ‘detected’, according to the ORForise platform. This meant that StORF-Reporter was only able to search for StORFs in the non-vitiated regions of the genome. For each model organism, between one and 26 non-vitiated missed Ensembl genes were recovered by using StORF-Reporter after the Prodigal annotation.

DIAMOND blastp was used to compare StORFs recovered from each model organism against the Swiss-Prot database and the proteomes derived from each respective model organism (see supplementary Table 6). There were StORFs that had a high sequence similarity to proteins in Swiss-Prot that were not present in either the Prodigal or Ensembl annotations, and were investigated further.

### StORF-Reporter finds complete CDS genes not present in Ensembl annotations

StORF-Reporter found many StORFs representing potential genes in the URs of the curated Ensembl annotations for each of the six model organisms. A number of StORFs had high sequence similarity to known protein coding genes in the Swiss-Prot protein database (see Table 2). Each of these StORFs have the potential to contain not only undiscovered genes but also historic gene fragments or other functional units waiting to be characterised. There were also a number of StORFs long enough to be genes which did not have a high level of sequence similarity to Swiss-Prot (see Supplementary Figure 4 for further detail on StORF lengths with and without Swiss-Prot hits). This could indicate that these sequences are not yet present in the databases or are fragmented in the genome.

**Table 2.** Table containing the number of StORFs found in the unannotated regions recovered from Ensembl annotations for six model organisms. The numbers of StORFs which had a high sequence similarity and >=80% subject coverage to a protein in Swiss-Prot and Ensembl proteome are listed.

| Model Organism | # URs/StORFs | Swiss-Prot [Coverage $\geq 80\%$] | Intra-Genome [Coverage $\geq 80\%$] |
| --- | --- | --- | --- |
| *B. subtilis* | 2,711 / 2,322 | 26 [20] | 8 [2] |
| *C. crescentus* | 2,321 / 1,554 | 8 [5] | 15 [3] |
| *E. coli* | 2,743 / 2,798 | 148 [107] | 57 [13] |
| *M. genitalium* | 157 / 168 | 63 [4] | 50 [0] |
| *P. fluorescens* | 3,509 / 3,404 | 73 [46] | 173[42] |
| *S. aureus* | 2,556 / 2,503 | 23 [4] | 17 [3] |

To study the possibility that StORFs may capture instances of gene-duplication, for each of the model organisms, the reported StORFs were aligned to the proteomes of each of the Ensembl model organisms. Between 8 and 173 StORFs were observed to contain an alignment to putative duplicate proteins from within the same genome. These StORFs are likely to represent duplicate genes that have not been detected by the current genome annotation methods or have undergone some level of deleterious mutation. While we observed 50 intra-genome StORFs ‘hits’ in *M. genitalium*, it was the only model organism not to contain a StORF with an alignment of >=80% of an Ensembl sequence. This is likely due to the ‘UGA/TGA’ codon repurposing. These results are reported in Table 2.

### Extending the E. *coli* pangenome

StORF-Reporter found StORFs that both extend the core-gene set and create novel core-gene clusters potentially expanding our understanding of the *E. coli* pangenome.

219 *E. coli* genomes were extracted from the collection of 6,223 Ensembl Bacterial genomes. From the 1,042,068 Ensembl protein sequences annotated in these genomes, 34,737 gene-clusters were formed, of which 20,676 were non-singletons. A median number of 3,038 URs and 2,958 StORFs were identified from the 219 genomes for a total of 673,136 URs and 652,056 StORFs (see Supplementary Table 7 for more detail). Representatives from each of the Ensembl-Only clusters (all 34,737 - including singletons) were combined with the 652,056 StORFs identified from the 219 *E. coli* genomes and the same CD-Hit clustering methodology was applied. This resulted in a total of 86,579 clusters, of which 31,676 were non-singletons. There were 51,929 StORF-Only clusters and 28,094 Ensembl-Only clusters. The remaining clusters contained both StORF and Ensembl sequences (Ensembl-StORF clusters).

The clustering of the Ensembl proteins resulted in 2,612 core-gene (found in >=99% of genomes) gene families.This is a similar figure to that of the number of core-genes found in other studies of *E. coli* strains (37). Most interestingly, with the addition of the StORF sequences, many Ensembl accessory clusters were extended into the soft-core and core (67 and 178 respectively) thresholds (see Table 3 for further detail). These constitute gene families which are likely to have been left out of many studies due to them incorrectly being classified as not part of the core. Additionally, there were clusters which contained both Ensembl and StORF sequences classified as core-gene families on the basis of their StORF sequences alone. While they were grouped with additional Ensembl genes, the Ensembl genes did not contribute to the distribution of the cluster (9 were core, 15 were soft-core and 648 were accessory). Most interestingly, there were 216 and 239 StORF-Only clusters with sequences from more than soft-core and core of the *E. coli* genomes, respectively. These clusters constitute entirely novel soft-core and core gene families missing from the Ensembl annotations. Lastly, there were 162 Ensembl clusters that although did cluster with StORF sequences, none of the StORFs were from additional genomes and as such were not counted in the above data.

**Table 3.** *Escherichia coli* pangenome gene families calculated from the set of 219 genomes. Definitions of the gene family category are as follows: Core-genes >=99%, Soft-core genes > = 95% to < 99% and Accessory genes >= 15% and < 95%. Gene families are only present in one category and Cluster Types are described in the methods.

| Cluster Types | Core | Soft-Core | Accessory |
| --- | --- | --- | --- |
| Ensembl-Only | 2,612 | 455 | 2,597 |
| Ensembl-StORF | 178 | 67 | 610 |
| StORF-Comb-Ensembl | 0 | 1 | 11 |
| StORF | 9 | 15 | 648 |
| StORF-Only | 239 | 216 | 3,426 |

Supplementary Figure 5 shows that the distribution of the unique *E. coli* genomes present in the Ensembl-Only, Ensembl-StORF and StORF-Only clusters all followed a similar U-shaped curve. Interestingly, the size of the clusters that were often extended by the StORFs was often increased by an substantial amount. For example, the proportion of clusters containing Ensembl sequences with 10 or fewer unique genomes was reduced from just below 60% to 40% with the addition of the StORF sequences. Only 87 of the 6,643 Ensembl gene families that clustered with StORF sequences did not extend into additional *E. coli* genomes.

The inherently limiting process of using hard-coded sequence similarity and length cut-offs to distinguish gene families has inevitably brought forward a number of interesting cases. One example can be seen in the combining of multiple Ensembl protein representative sequences, and thus gene families, into a single new cluster by the addition of StORFs. This was a result of StORF sequences effectively bridging the gap between multiple Ensembl sequences (defined henceforth as StORF-Combined-Ensembl clusters). The 71 amino acid representative protein AKD71933 of Ensembl cluster 4,714 (consisting of 59 sequences and spanning 24 strains, some of which have duplicates within the same genome), was clustered with the representative 66 amino acid protein AHM40952 of Ensembl cluster 6,112 (consisting of 26 sequences from 26 strains) when StORF sequences were included in the second round of CD-Hit clustering. Previously, these two Ensembl proteins, reported as separate transposases in NCBI’s non-redundant protein database (38), were distinct enough to be clustered separately. However, they were both clustered together in combined Cluster 4 with 335 StORFs from 99 strains. The representative StORF sequence from this cluster was identified to be a ‘Insertion element protein’ in the same database. The StORFs in this cluster are longer than the Ensembl genes (around 100 aa), and so CD-Hit chose a StORF to be its representative sequence for the combined cluster. As can be seen in the Clustal Omega (39) multiple sequence alignment of the representative sequences of the above-mentioned clusters in Figure 4, their sequence identity was high, ≥ 90% (the minimum CD-Hit percentage identity for clustering) along the regions in which they aligned.

**Figure 4.**
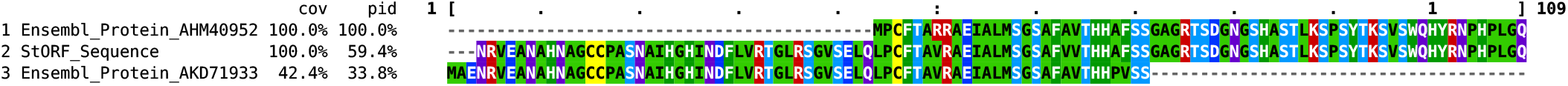
Clustal Omega multiple sequence alignment of the two Ensembl representative sequences, AHM40952 and AKD71933, which were combined in *E. coli* pangenome cluster 4 by the additional StORF sequence. This combination is done because the StORF sequence extends to where AKD71933 begins and to where AKD71933 ends. These two sequences on their own do not align together for more than a small portion of either of their full sequence length, thus the reason they formed independent clusters.

A further example of this type of clustering can be seen in Supplementary Figure 6 which reports a multiple sequence alignment of three Ensembl representative sequences (AHM40736, AIT36070 and AFS84250) combined into a new cluster with a single StORF sequence (Cluster 17,575). As we can observe from their alignment, they are very similar and are further examples of the limitations we impose when attempting to categorise natural elements into computationally-friendly representations. There were 77 of these StORF-Combined-Ensembl clusters which together had clustered 164 Ensembl representative sequences together. These clusters are presented separately in Table 3.

### StORFs identified within and across multiple genera

The previous analyses of URs and StORFs have been conducted on the same species. It could therefore, be assumed that many of the StORFs which have been identified are remnants from a multitude of different genomic factors such as assembly error, genomic structural elements or additional processes we are yet to fully understand. However, in this study, large numbers of StORFs were identified with and without similarity to Ensembl genes across multiple genera, forming both novel and extended cross-genera gene families.

The complete collection of 21,503,164 Ensembl-annotated protein sequences was collected from the 6,223 Ensembl Bacteria genomes (see Supplementary Table 1 for details on the genomes used). A median number of 2,305 and 1,981 URs and StORFs were identified for each of the 6,223 genomes for a total of 14,221,482 and 13,301,175 sequences respectively (see Supplementary Table 8 for more detail). Using CD-Hit, Ensembl-Only, Ensembl-StORF, and StORF-Only clusters were identified spanning multiple genera across the 179 genera from Ensembl Bacteria. The distribution of genera in the Ensembl-Only clusters is wide, and many clusters are formed from more than ten different genera (see Table 4). Furthermore, not only have Ensembl-Only clusters been added to and extended by StORFs (creating Ensembl-StORF clusters), but also substantial numbers of StORF-Only clusters have been found with StORFs from multiple genera. Furthermore, some clusters were identified with Ensembl and StORF sequences from the same genera and genomes. However, although these are likely to represent important candidates for gene duplication studies and should be added to canonical annotations, they were not the aim of this study.

**Table 4.** The number of clusters which have sequences from multiple genera. The five cluster types are; (1) Ensembl-Only, (2) Ensembl-StORF, which are the clusters which have been extended into their respective genera group by the addition of StORF sequences, (3) ‘StORF-Combined-Ensembl’ (StORF-Comb-Ensembl), which reports the number of gene families where StORF sequences combined at least 2 or more Ensembl cluster representatives together, (4) StORF, which are the same clusters as Ensembl-StORF but are counted only by their number of StORF sequences and (5) StORF-Only, which are the clusters which only contain StORF sequences and thus did not cluster with any Ensembl sequence. StORF-Only clusters with a single sequence were not included in these results.

| Cluster type | 1 Genus | 2 Genera | 3 Genera | 4 Genera | 5 Genera | 6 Genera | >6 Genera |
| --- | --- | --- | --- | --- | --- | --- | --- |
| Ensembl-Only | 1,569,403 | 6,9884 | 6,323 | 2,491 | 1,338 | 803 | 1,427 |
| Ensembl-StORF | 0 | 0 | 142 | 69 | 38 | 19 | 37 |
| StORF-Comb-Ensembl | 0 | 0 | 0 | 1 | 0 | 0 | 5 |
| StORF | 125,357 | 2,530 | 247 | 63 | 29 | 10 | 25 |
| StORF-Only | 1,087,592 | 27,079 | 2,313 | 813 | 367 | 172 | 265 |

StORFs have changed the diversity and distribution of gene families by extending the canonical Ensembl gene clusters into additional genera which were not annotated in the Ensembl annotations. One example of this, is a singleton Ensembl-Only cluster (Cluster 1,845,585) that consisted of a 35 amino acid *E. coli* Ensembl gene (reported in GenBank (40) as a hypothetical protein - *E. coli kte42* - gene ELF73632). This sequence, when combined with the StORF sequences from all 6,223 Ensembl genomes, was clustered into the 4*^th^* largest combined cluster (Ensembl-StORF combined ‘Cluster 3’), with StORFs spanning multiple different genera. This cluster consisted of 1,162 StORF sequences from a total of 149 genomes and 5 genera (*Shigella, Escherichia, Pantoea, Klebsiella* and *Enterobacter*). The fact that the only organism to have this sequence annotated was *E. coli* may not be a coincidence, *E.coli* being the most studied model organism.

As another example, the small Ensembl Cluster 83,472, which consisted of 32 sequences between 63-70 amino acids long and spanning 4 genera (*Escherichia, Streptococcus, Yersinia* and *Shigella*) was significantly extended by 726 StORF sequences from 7 additional genera (*Pantoea, Klebsiella, Citrobacter, Enterobacter, Serratia, Salmonella* and *Bacillus*) as part of the Ensembl-StORF combined Cluster 14. The function of this protein is unclear. An alignment to the NCBI non-redundant protein database reported a number of different annotations which indicate that it could be either part of a transposase family, a plasmid stability protein or something else entirely.

As with the *E. coli* pangenome study, the hard-coded cutoffs used for the CD-Hit gene family analysis produced a number of clusters which combined multiple Ensembl representative sequences which were previously clustered separately. Six Ensembl-StORF combined clusters were found to contain more than one Ensembl representative sequence and together combined 14 Ensembl-Only clusters. Additionally, many of the StORF sequences were found in genera that were not present in the original and separate Ensembl-Only clusters. An example of one such cluster, Ensembl-StORF cluster 124,470, combined three Ensembl-Only representatives together with seven StORF sequences. Supplementary Figure 7 shows the phylogenetic tree built with ClustalO and FastTree (41) and plotted with iTOL (42) (see Supplementary Figure 8 for the multiple sequence alignment). This tree clearly reports the diversity and, at the same time, the similarity of the StORFs and Ensembl annotated sequences. Additionally, the multiple sequence alignment reports how for some amino acid positions, StORF sequences can be more similar to one or more Ensembl gene as they are compared to each other. Therefore, it could be argued that as these StORF sequences bridge the gap between the three separate Ensembl gene families, they have changed the dynamics of these inter-genera gene families. These type of clusters were included in the results of Table 4, separately in row ‘StORF-Comb-Ensembl’.

StORF-Reporter was also able to identify a number of clusters containing only StORF sequences that spanned many different genera. The biggest of these, combined cluster 15,543, spans 22 genera and consists of 81 genomes and 92 sequences. At 380 amino acids long, the representative of this cluster had a 90% percent identity and 55% query coverage to the ‘fatty-acid-CoA ligase fadD18’ protein in GenBank. This large, widespread protein was mostly observed as a single gene in each genome but in 5 species it was found duplicated with between 2 and 6 copies in non-contiguous URs.

## DISCUSSION

### The unannotated regions of prokaryote genomes contain many potential genes

The high gene densities observed in canonical prokaryotic annotations have long been “perceived as evidence of adaptive genome streamlining” (1). However, this is in contradiction with the observations that many prokaryote genomes contain a large number of long URs that comprise between 10-20% of their genomes (see Table 2 and Supplementary Tables 3 and 4). Despite all advancements, there is still scope to improve genome annotation strategies. Previous findings (2) confirmed that contemporary methods such as Prodigal (33) accurately detect the majority of CDS genes (in genomes which use the ‘universal’ codon table). However, as shown in this work, most genomes exhibited thousands of URs that contain genes missed by state-of-the-art gene prediction methods and are thus missing from the canonical Ensembl annotations or exhibited high levels of sequence identity to proteins in the Swiss-Prot database.

It is difficult to know which proportion of the StORFs identified in this study are a result of the current ineffectiveness of genome annotation methods or are in fact genes which are no longer expressed, possibly due to mutations in a promoter region (putative pseudogenes). This is a problem that is not likely to be solved soon, as evidenced by the large numbers of “hypothetical” or “putative” genes that exist in canonical genome annotations, even in model organisms like *E. coli* (43). Interestingly, the stop codon abundances and usages presented in Supplementary Tables 15 and 16, suggest that StORFs without homology to known CDS genes may be at least partially validated with stop codon usage analysis. Additionally, since the coding and non-coding regions of prokaryote genomes are believed to evolve separately, reacting to intra- and inter-genomic pressures independently (44), SNP and mutation rate studies could also be potentially used as evidence that they are functional genes. In addition, the observation that many of the StORFs appear to be gene duplicates and/or members of broad gene families, validates that our approach can identify real genes (see ‘Intra-Genome’ results in Table 2). Regardless of whether these duplicates are pseudogenes, a type of gene that is still too often ignored, their presence is important and their addition to prokaryotic annotations is likely to enhance our understanding of how these genomes evolve. For example, we have identified large numbers of StORF-Only clusters, those gene families consisting only of StORF sequences that did not have a high level of similarity to any known genes, spanning over 10 genera in some cases. These may represent CDS sequences or cryptic genomic elements, previously overlooked, and are prime targets for future investigations.

### The ‘Stop - Open Reading Frame’ is a valuable and unambiguous concept

As previously mentioned, our definition of a StORF is synonymous with the Sequence Ontology (9) definition of an ORF - “The in-frame interval between the stop codons of a reading frame.”. This differs with the canonical use of the term ‘ORF’ (a coding frame beginning with a start codon and ending with a stop codon) which has become the standard approach for most genome annotation tools (2). Unlike the use of ‘ORF’ in genome annotation, StORFs have a unambiguous description and use. Additionally, we have shown the value of StORFs in their unbiased identification of genes, even those with non-canonical start codon profiles. Lastly, as StORFs are only reported from the URs already identified in previously annotated genomes, we are building upon the excellent work of previous researchers, rather than restarting annotations from scratch. The StORF_Adder tool is provided with the ORForise package (2), to supplement existing genome annotations from Ensembl or workflows such as PROKKA, retaining the same output format.This allows downstream analysis tools such as Roary (45) and Coinfinder (46) to interpret these new annotations without further intervention.

### StORFs can extend pangenomes

For some time, it has been known that a single genome is not enough to characterise the functional profile of a species requiring instead a pangenomic approach. As pangenomes consist of a species-wide collection of core-genes that are present in most/all genomes and accessory genes that are only present in some genomes, missing annotations may impact our interpretation of whether certain gene families are core or not. Indeed, while our pangenomic study of *E. coli* has shed light on the potential for yet more diversity in its accessory gene collection, it is the much increased set of core-genes, both expanded and novel which truly showcase the utility of StORF-Reporter. These gene families are candidates for studying how a species pangenome can adapt to ever-changing environments (47) as they have been retained against the pressure for genome streamlining (48, 49, 50) and are more likely to be functionally important. Fundamentally, while the identification of these StORFs has provided additional genomic knowledge previously left in the dark corners of the URs of their genomes, to truly use the nascent information they may harbour, much expanded experimental work is needed.

### StORFs can extend intra- and inter-genera gene collections

The results of the StORFs reported in the six model organisms and across the *E. coli* pangenome have shown wide levels of diversity and spread. However, it could be possible that many of these StORFs are artefacts of genome assembly and are species-specific structural elements and not primarily CDS genes. Therefore, the inter-genera analysis of the 6,223 genomes from Ensembl Bacteria was key not only to study the presence of StORFs in other diverse species, but also to provide evidence to further validate them as putative CDS genes.

StORFs were found across many different genera at high sequence similarities (>=95%) (see Table 4). As with the *E. coli* pangenomic study, many StORFs from the same cluster were found in different locations across the various genomes, or in the case of duplication, in different regions and thus URs of the same genome. The presence of these StORFs, adds credence to the hypothesis that they are not structural elements or other genomic artefacts but instead functional (past or present) CDS genes.

As StORF-Reporter is species- and start codon-agnostic, we are better able to observe the full spectrum of variation within a gene family. This provides yet further justification of the need for species-agnostic gene prediction methodologies which do not rely on established model organisms or databases.

### Are some StORFs pseudogenised genes?

The potential diversity of all possible protein sequences is vast (51). A typical protein length of about 300 amino acids can have theoretically up to 20^300^ different possible polypeptide chains, yet current protein databases only contain a tiny fraction of this, such as Swiss-Prot, which contains a little over 500,000 protein sequences. Many of these are duplicates or very similar to each other and many of these are not experimentally validated. Adding to this complexity is the vast number of pseudogenes, which were likely functional proteins in the recent past, identified through the extensive genome sequencing of the last few decades. While mutation rates have long been studied in prokaryotes (52, 53), there are still unresolved questions surrounding the speed at which pseudogenes are created and subsequently removed from the genome. Some studies have shown that pseudogenes are more likely to be a result of failed horizontal gene transfer (54). Furthermore, while archaea and most non-pathogenic bacteria exhibit greater retention of ancient gene remnants, obligate pathogenic bacteria tend to have younger pools of pseudogenes (54). As such, the types of genes pseudogenised and the rate at which they are mutated are highly variable between species and are likely to be related to environmental specific cues.

Unfortunately, current methods fail to address the prevalence of gene pseudogenisation; therefore, the true extent of gene pseudogenisation is likely to remain unclear for the foreseeable future. As reported by Goodhead *et al* (55), of the five different process causing a gene to be pseudogenised (premature stop, loss/change of function domain, loss of start codon, loss of promoter/RBS, frameshift), a StORF has the potential to identify all of them (some in fragmentary form). Additionally, while a single StORF could only report one fragment of a gene pseudogenised by a premature stop codon, the combination of multiple consecutive-StORFs in the same reading frame has the potential to recover the entire sequence of a pseudogenised gene.Furthermore, as a StORF captures a portion of the 5’ untranslated region of a CDS gene, it may also identify gene silencing mutations, such as start codon or ribosomal binding site mutations.

## CONCLUSION

Genome annotation is a mature but continually developing field. With each new sequencing project, the proportion of uncharacterised sequences found can vary significantly, and often hundreds of novel genes are discovered with varying similarity to those from previously sequenced taxa (56), suggesting that considerable gene diversity remains to be discovered. Historical and systematic biases and errors mean that some types of genes will almost always be missed in these studies, resulting in many tools reporting the same URs (2). This might lead researchers to feel a sense of false security when using canonical annotation tools, but, as we have demonstrated, the StORF-Reporter methodology can enhance current annotations, reporting many missed genes in these regions.

One major limiting factor shared across many if not all genome annotation analysis studies is the use of a genomic database such as Ensembl Bacteria as a ground truth (under the acceptance that they represent a complete reference). In this study, we tested that assumption of completeness. We used StORF-Reporter to investigate the URs extracted from over 6,000 Ensembl annotated genomes. StORF-Reporter was able to identify more than 200 StORF-Only core-gene families from the *E. coli* pangenome, which were distinct from any of those present in Ensembl. This constitutes a near 10% increase in the size of the core-gene collection. Moreover, a large number of StORFs were highly similar to known proteins present across multiple genera. Both sets of StORFs showcase how even highly-conserved gene families are still missing from canonical annotations. This is strong evidence that StORF-Reporter is able to recover novel functional sequences even when applied to well-studied reference genomes from model organisms.

The URs and StORFs identified by StORF-Reporter can be interpreted and used in a number of different ways. Once StORFs have been recovered, the remaining regions of DNA in the collection of URs for any genome are not without significance. The use of StORF-Reporter makes it easier to study these cryptic regions, which could be classified as the ‘real unknown-unknowns’. We provide a tool for the extraction of these unknown-unknowns in the StORF-Reporter package named “Un-StORFed”.

In summary, genomes are not static; elements are added, removed, and mutated over time, and thus the content of a genome is dynamic in nature. This work represents a movement toward the thesis that every nucleotide in the genome should be labelled, regardless of whether it belongs to a functional gene, a pseudogene, a gene fragment that is no longer functional or any other genomic feature. Therefore, we encourage the community to use StORF-Reporter to enhance both canonical and novel genome annotations.

## DATA AVAILABILITY

The StORF-Reporter package is available on (https://github.com/NickJD/StORF-Reporter) and as a PyPi package (https://pypi.org/project/StORF-Reporter/). The full ORForise package which has been updated with the StORForise plugin is also available on Github (https://github.com/NickJD/ORForise) and as a PyPi package (https://pypi.org/project/ORForise/). The extracted URs and StORFs from the entire collection of 6,233 Ensembl genomes is available at the Open Science Framework (DOI: 10.17605/OSF.IO/4DWTN).

## AUTHOR CONTRIBUTIONS

All authors discussed the conceptualisation of StORFs and their impact. N.J.D wrote the code. All authors contributed to the manuscript.

## REFERENCES

1. Itamar Sela, Yuri I Wolf, and Eugene V Koonin. Theory of prokaryotic genome evolution. Proceedings of the National Academy of Sciences, 113(41):11399–11407, 2016.

2. Nicholas J Dimonaco, Wayne Aubrey, Kim Kenobi, Amanda Clare, and Christopher J Creevey. No one tool to rule them all: Prokaryotic gene prediction tool annotations are highly dependent on the organism of study. Bioinformatics, 38(5):1198–1207, 2021.

3. Ryan J Taft, Michael Pheasant, and John S Mattick. The relationship between non-protein-coding DNA and eukaryotic complexity. Bioessays, 29(3):288–299, 2007.

4. Matthew R Hemm, Brian J Paul, Thomas D Schneider, Gisela Storz, and Kenneth E Rudd. Small membrane proteins found by comparative genomics and ribosome binding site models. Molecular Microbiology, 70(6):1487–1501, 2008.

5. Jayavel Sridhar, Radhakrishnan Sabarinathan, Shanmugam Siva Balan, Ziauddin Ahamed Rafi, Paramasamy Gunasekaran, and Kanagaraj Sekar. Junker: an intergenic explorer for bacterial genomes. Genomics, Proteomics & Bioinformatics, 9(4-5):179–182, 2011.

6. Chen-Hsun Tsai, Rick Liao, Brendan Chou, Michael Palumbo, and Lydia M Contreras. Genome-wide analyses in bacteria show small-RNA enrichment for long and conserved intergenic regions. Journal of Bacteriology, 197(1):40–50, 2015.

7. Harry A Thorpe, Sion C Bayliss, Laurence D Hurst, and Edward J Feil. Comparative analyses of selection operating on nontranslated intergenic regions of diverse bacterial species. Genetics, 206(1):363–376, 2017.

8. Patricia Sieber, Matthias Platzer, and Stefan Schuster. The definition of open reading frame revisited. Trends in Genetics, 34(3):167–170, 2018.

9. Karen Eilbeck, Suzanna E Lewis, Christopher J Mungall, Mark Yandell, Lincoln Stein, Richard Durbin, and Michael Ashburner. The Sequence Ontology: a tool for the unification of genome annotations. Genome Biology, 6(5):R44, 2005.

10. L Dalgarno and J Shine. Conserved terminal sequence in 18s rRNA may represent terminator anticodons. Nature New Biology, 245(148):261–262, 1973.

11. Douglas F Browning and Stephen JW Busby. The regulation of bacterial transcription initiation. Nature Reviews Microbiology, 2(1):57–65, 2004.

12. Thomas Dandekar, Berend Snel, Martijn Huynen, and Peer Bork. Conservation of gene order: a fingerprint of proteins that physically interact. Trends in Biochemical Sciences, 23(9):324–328, 1998.

13. Andre Villegas and Andrew M Kropinski. An analysis of initiation codon utilization in the Domain Bacteria–concerns about the quality of bacterial genome annotation. Microbiology, 154(9):2559–2661, 2008.

14. Frida Belinky, Igor B Rogozin, and Eugene V Koonin. Selection on start codons in prokaryotes and potential compensatory nucleotide substitutions. Scientific Reports, 7(1):1–10, 2017.

15. Pavel V Baranov, John F Atkins, and Martina M Yordanova. Augmented genetic decoding: global, local and temporal alterations of decoding processes and codon meaning. Nature Reviews Genetics, 16(9):517–529, 2015.

16. Frida Belinky, Vladimir N Babenko, Igor B Rogozin, and Eugene V Koonin. Purifying and positive selection in the evolution of stop codons. Scientific Reports, 8(1):1–11, 2018.

17. Inna S Povolotskaya, Fyodor A Kondrashov, Alice Ledda, and Peter K Vlasov. Stop codons in bacteria are not selectively equivalent. Biology Direct, 7(1):1–13, 2012.

18. Herman Tse, James J Cai, Hoi-Wah Tsoi, Esther PT Lam, and Kwok-Yung Yuen. Natural selection retains overrepresented out-of-frame stop codons against frameshift peptides in prokaryotes. BMC Genomics, 11(1):1–13, 2010.

19. Torsten Seemann. Prokka: rapid prokaryotic genome annotation. Bioinformatics, 30(14):2068–2069, 2014.

20. Kevin L Howe, Bruno Contreras-Moreira, Nishadi De Silva, Gareth Maslen, Wasiu Akanni, James Allen, Jorge Alvarez-Jarreta, Matthieu Barba, Dan M Bolser, Lahcen Cambell, et al. Ensembl Genomes 2020 – enabling non-vertebrate genomic research. Nucleic Acids Research, 48(D1):D689–D695, 2020.

21. Guido Van Rossum and Fred L. Drake. Python 3 Reference Manual. CreateSpace, Scotts Valley, CA, 2009.

22. Niv Sabath, Dan Graur, and Giddy Landan. Same-strand overlapping genes in bacteria: compositional determinants of phase bias. Biology Direct, 3(1):36, 2008.

23. Alexandra M Schnoes, Shoshana D Brown, Igor Dodevski, and Patricia C Babbitt. Annotation error in public databases: misannotation of molecular function in enzyme superfamilies. PLoS Computational Biology, 5(12):e1000605, 2009.

24. Andrew S. Warren, Jeremy Archuleta, Wu-chun Feng, and Joao Carlos Setubal. Missing genes in the annotation of prokaryotic genomes. BMC Bioinformatics, 11(1):131, 2010.

25. Derrick E Wood, Henry Lin, Ami Levy-Moonshine, Rajiswari Swaminathan, Yi-Chien Chang, Brian P Anton, Lais Osmani, Martin Steffen, Simon Kasif, and Steven L Salzberg. Thousands of missed genes found in bacterial genomes and their analysis with COMBREX. Biology Direct, 7(1):37, 2012.

26. Tit-Yee Wong, Sanjit Fernandes, Naby Sankhon, Patrick P Leong, Jimmy Kuo, and Jong-Kang Liu. Role of premature stop codons in bacterial evolution. Journal of Bacteriology, 190(20):6718–6725, 2008.

27. Natalia N Ivanova, Patrick Schwientek, H James Tripp, Christian Rinke, Amrita Pati, Marcel Huntemann, Axel Visel, Tanja Woyke, Nikos C Kyrpides, and Edward M Rubin. Stop codon reassignments in the wild. Science, 344(6186):909–913, 2014.

28. Kevin Dybvig and LeRoy L Voelker. Molecular biology of Mycoplasmas. Annual Reviews in Microbiology, 50(1):25–57, 1996.

29. Alexey V Lobanov, Anton A Turanov, Dolph L Hatfield, and Vadim N Gladyshev. Dual functions of codons in the genetic code. Critical Reviews in Biochemistry and Molecular Biology, 45(4):257–265, 2010.

30. Thressa C Stadtman. Selenocysteine. Annual Review of Biochemistry, 65(1):83–100, 1996.

31. Gayathri Srinivasan, Carey M James, and Joseph A Krzycki. Pyrrolysine encoded by UAG in archaea: charging of a UAG-decoding specialized tRNA. Science, 296(5572):1459–1462, 2002.

32. Fred Sherman, John W Stewart, and Susumu Tsunasawa. Methionine or not methionine at the beginning of a protein. Bioessays, 3(1):27–31, 1985.

33. Doug Hyatt, Gwo-Liang Chen, Philip F LoCascio, Miriam L Land, Frank W Larimer, and Loren J Hauser. Prodigal: prokaryotic gene recognition and translation initiation site identification. BMC Bioinformatics, 11(1):119, 2010.

34. Limin Fu, Beifang Niu, Zhengwei Zhu, Sitao Wu, and Weizhong Li. Cd-hit: accelerated for clustering the next-generation sequencing data. Bioinformatics, 28(23):3150–3152, 2012.

35. Benjamin Buchfink, Chao Xie, and Daniel H Huson. Fast and sensitive protein alignment using DIAMOND. Nature Methods, 12(1):59–60, 2015.

36. UniProt Consortium.UniProt: a worldwide hub of protein knowledge. Nucleic Acids Research, 47(D1):D506–D515, 2019.

37. Rohan Maddamsetti, Philip J Hatcher, Anna G Green, Barry L Williams, Debora S Marks, and Richard E Lenski. Core genes evolve rapidly in the long-term evolution experiment with Escherichia coli. Genome Biology and Evolution, 9(4):1072–1083, 2017.

38. Daniel H Haft, Michael DiCuccio, Azat Badretdin, Vyacheslav Brover, Vyacheslav Chetvernin, Kathleen O’Neill, Wenjun Li, Farideh Chitsaz, Myra K Derbyshire, Noreen R Gonzales, et al. RefSeq: an update on prokaryotic genome annotation and curation. Nucleic Acids Research, 46(D1):D851–D860, 2018.

39. Fabian Sievers and Desmond G Higgins. Clustal Omega for making accurate alignments of many protein sequences. Protein Science, 27(1):135–145, 2018.

40. Karen Clark, Ilene Karsch-Mizrachi, David J Lipman, James Ostell, and Eric W Sayers. Genbank. Nucleic Acids Research, 44(D1):D67–D72, 2016.

41. Morgan N Price, Paramvir S Dehal, and Adam P Arkin. FastTree 2– approximately maximum-likelihood trees for large alignments. PloS One, 5(3), 2010.

42. Ivica Letunic and Peer Bork. Interactive Tree Of Life (iTOL) v5: an online tool for phylogenetic tree display and annotation. Nucleic Acids Researchy, 49(W1):W293–W296, 2021.

43. Sankha Ghatak, Zachary A King, Anand Sastry, and Bernhard O Palsson. The y-ome defines the 35% of Escherichia coli genes that lack experimental evidence of function. Nucleic Acids Research, 47(5):2446–2454, 2019.

44. Igor B Rogozin, Kira S Makarova, Darren A Natale, Alexey N Spiridonov, Roman L Tatusov, Yuri I Wolf, Jodie Yin, and Eugene V Koonin. Congruent evolution of different classes of non-coding DNA in prokaryotic genomes. Nucleic Acids Research, 30(19):4264–4271, 2002.

45. Andrew J Page, Carla A Cummins, Martin Hunt, Vanessa K Wong, Sandra Reuter, Matthew TG Holden, Maria Fookes, Daniel Falush, Jacqueline A Keane, and Julian Parkhill. Roary: rapid large-scale prokaryote pan genome analysis. Bioinformatics, 31(22):3691–3693, 2015.

46. Fiona Jane Whelan, Martin Rusilowicz, and James Oscar McInerney. Coinfinder: detecting significant associations and dissociations in pangenomes. Microbial Genomics, 6(3), 2020.

47. David A Rasko, MJ Rosovitz, Garry SA Myers, Emmanuel F Mongodin, W Florian Fricke, Pawel Gajer, Jonathan Crabtree, Mohammed Sebaihia, Nicholas R Thomson, Roy Chaudhuri, et al. The pangenome structure of Escherichia coli: comparative genomic analysis of E. coli commensal and pathogenic isolates. Journal of Bacteriology, 190(20):6881–6893, 2008.

48. Susumu Ohno. Evolution by gene duplication. Springer Science & Business Media, 2013.

49. Anthony Levasseur and Pierre Pontarotti. The role of duplications in the evolution of genomes highlights the need for evolutionary-based approaches in comparative genomics. Biology Direct, 6(1):1–12, 2011.

50. Stephen J Giovannoni, J Cameron Thrash, and Ben Temperton. Implications of streamlining theory for microbial ecology. The ISME Journal, 8(8):1553–1565, 2014.

51. Bruce Alberts, Alexander Johnson, Julian Lewis, Martin Raff, Keith Roberts, and Peter Walter. The shape and structure of proteins. In Molecular Biology of the Cell. 4th edition. Garland Science, 2002.

52. Salvador E Luria and Max Delbrück. Mutations of bacteria from virus sensitivity to virus resistance. Genetics, 28(6):491, 1943.

53. William A Rosche and Patricia L Foster. Determining mutation rates in bacterial populations. Methods, 20(1):4–17, 2000.

54. Yang Liu, Paul M Harrison, Victor Kunin, and Mark Gerstein. Comprehensive analysis of pseudogenes in prokaryotes: widespread gene decay and failure of putative horizontally transferred genes. Genome Biology, 5(9):1–11, 2004.

55. Ian Goodhead and Alistair C Darby. Taking the pseudo out of pseudogenes. Current Opinion in Microbiology, 23:102–109, 2015.

56. Mark Kowarsky, Joan Camunas-Soler, Michael Kertesz, Iwijn De Vlaminck, Winston Koh, Wenying Pan, Lance Martin, Norma F Neff, Jennifer Okamoto, Ronald J Wong, et al. Numerous uncharacterized and highly divergent microbes which colonize humans are revealed by circulating cell-free DNA. Proceedings of the National Academy of Sciences, 114(36):9623–9628, 2017.

