## Supplemental Document for "StORF-Reporter: Finding Genes between Genes"

### Supplementary Information
### StORF-Reporter: Finding Genes between Genes

Nicholas J. Dimonaco [1,2*], Wayne Aubrey [2], Kim Kenobi [3],
Amanda Clare [2]†, and Christopher J. Creevey [4]†

[1]Institute of Biological, Environmental and Rural Sciences, Aberystwyth University, Aberystwyth, SY23 3PD, Wales, UK
[2]Department of Computer Science, Aberystwyth University, Aberystwyth, SY23 3PD, Wales, UK
[3]Department of Maths and Physics, Aberystwyth University, Aberystwyth, SY23 3PD, Wales, UK
[4]School of Biological Sciences, Queen's University Belfast, Belfast, BT7 1NN, Northern Ireland, UK

†The authors wish it to be known that, in their opinion, the last two authors should be regarded as Joint Senior Authors.

### 1 Supplementary Figures

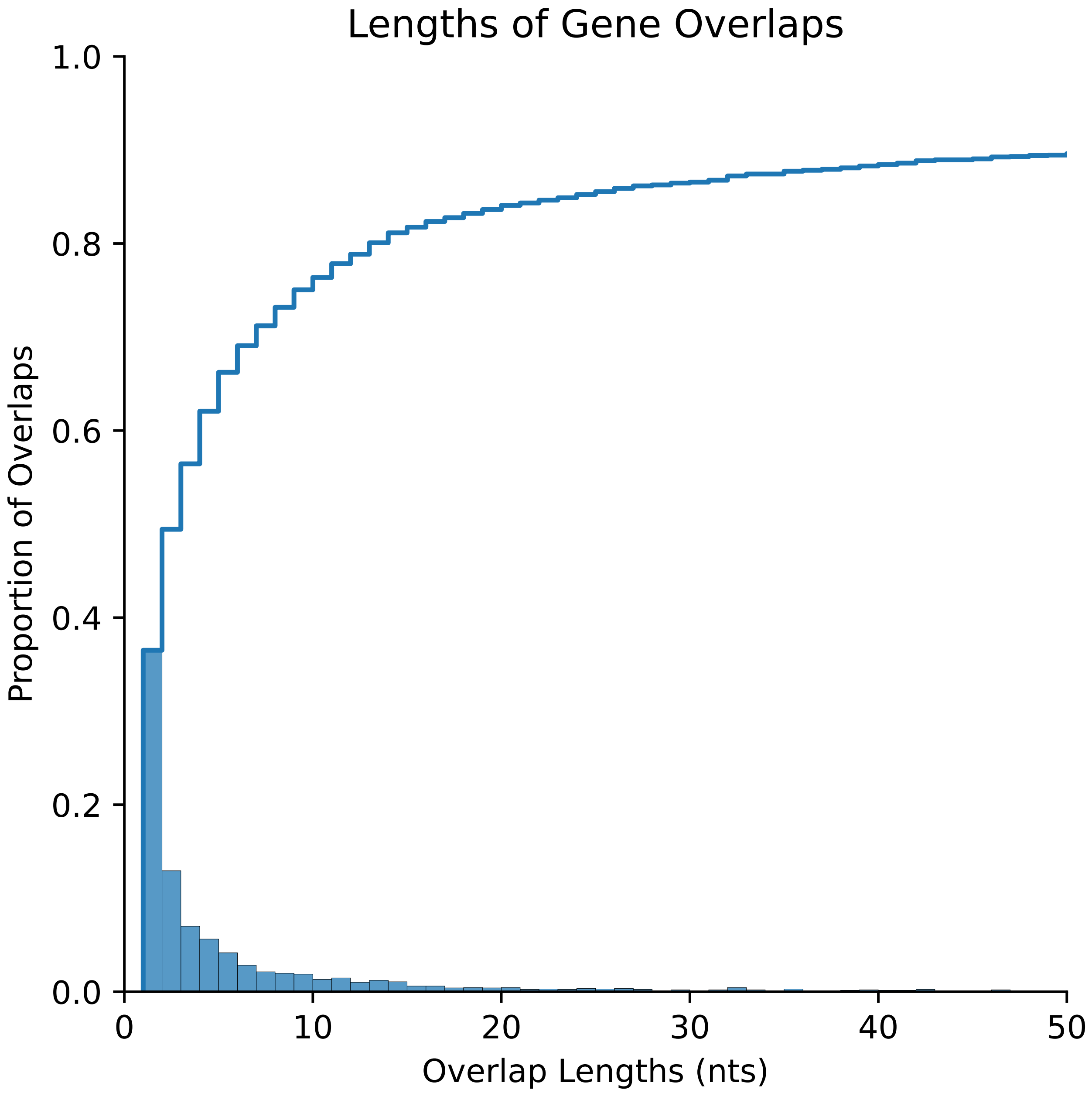


Figure 1: Shown here are the proportional overlap lengths in nucleotides between all CDS genes from the 6,223 filtered genomes from Ensembl Bacteria. The blue line reports the cumulative proportion of gene overlaps increasing very little after 10-20 nt.

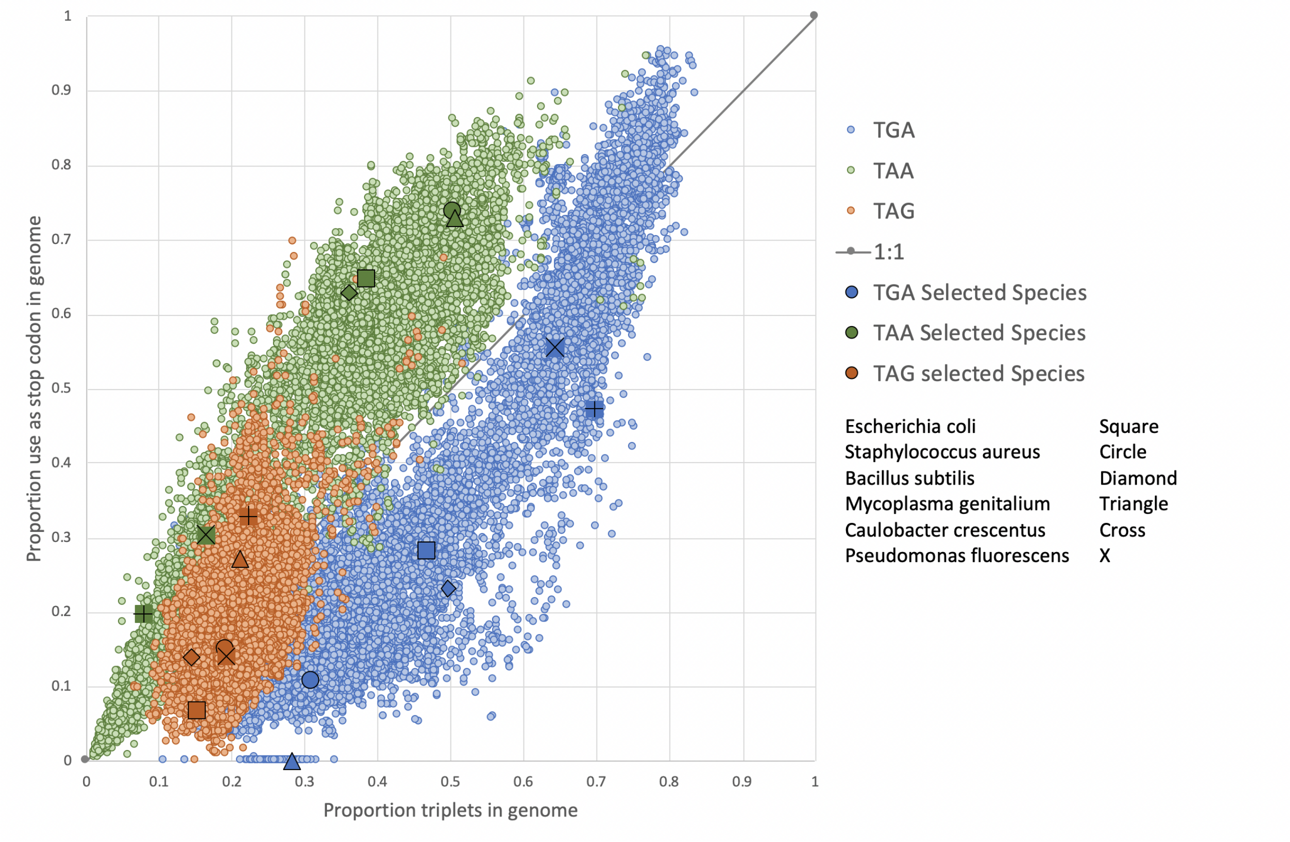


Figure 2: Graph showing stop codon usage (y axis) and proportion of triplets (x axis) in the 44,048 genomes from Ensembl Bacteria. For each of the 6 model organisms, this show the distribution across the entire genome of the three triplets TAA, TAG and TGA in all six reading frames and stop codon usage (i.e. the actual relative usage in Ensembl CDS genes of the three different stop codons).

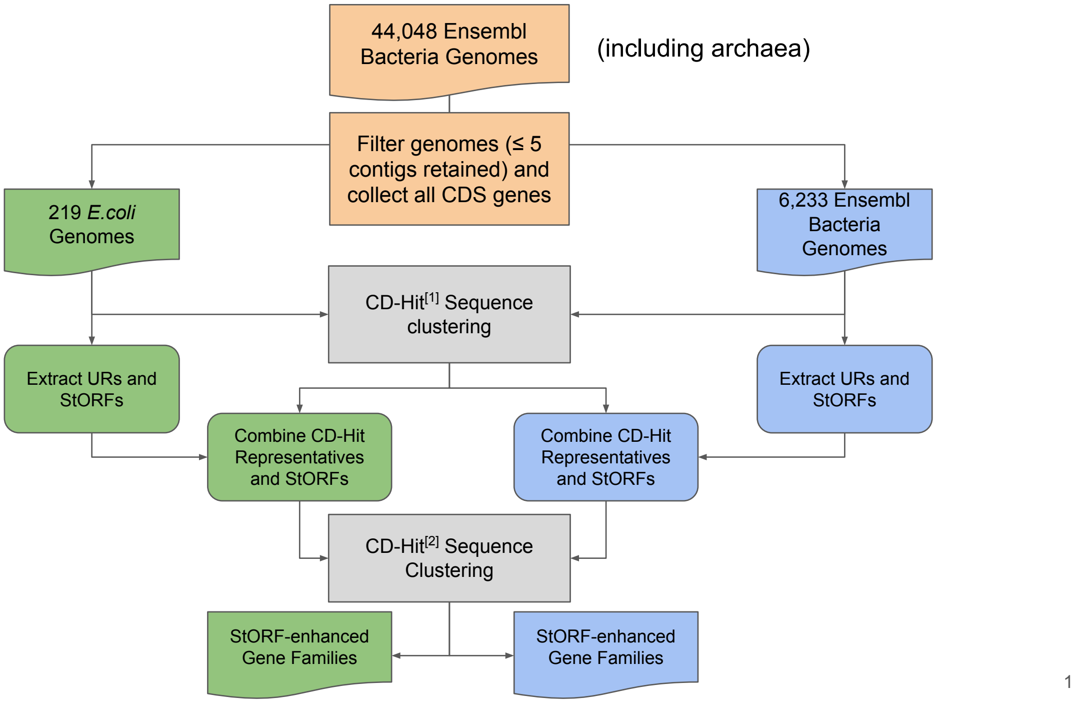


Figure 3: This workflow diagram presents the process used to create the *E. Coli* pangenomic and cross-genera gene families. The same original Ensembl input data and genome filtering was used for both studies (orange boxes) and the same CD-Hit clustering protocol was undertaken (grey boxes). The green boxes indicate the specific route the *E. coli* genomes took and the blue boxes report the same for the cross-genera study. There are two separate CD-Hit stages which are applied the same to both datasets and are described as :[1] This CD-Hit clustering stage was performed only on the amino acid sequences reported in the Ensembl Bacteria annotations, [2] This CD-Hit clustering stage was performed on the Ensembl amino acid sequence representatives reported by the previous CD-Hit analysis and the StORFs identified from the Ensembl annotations. The same parameters of 90% sequence identity and shorter sequence length cut offs were applied to both.

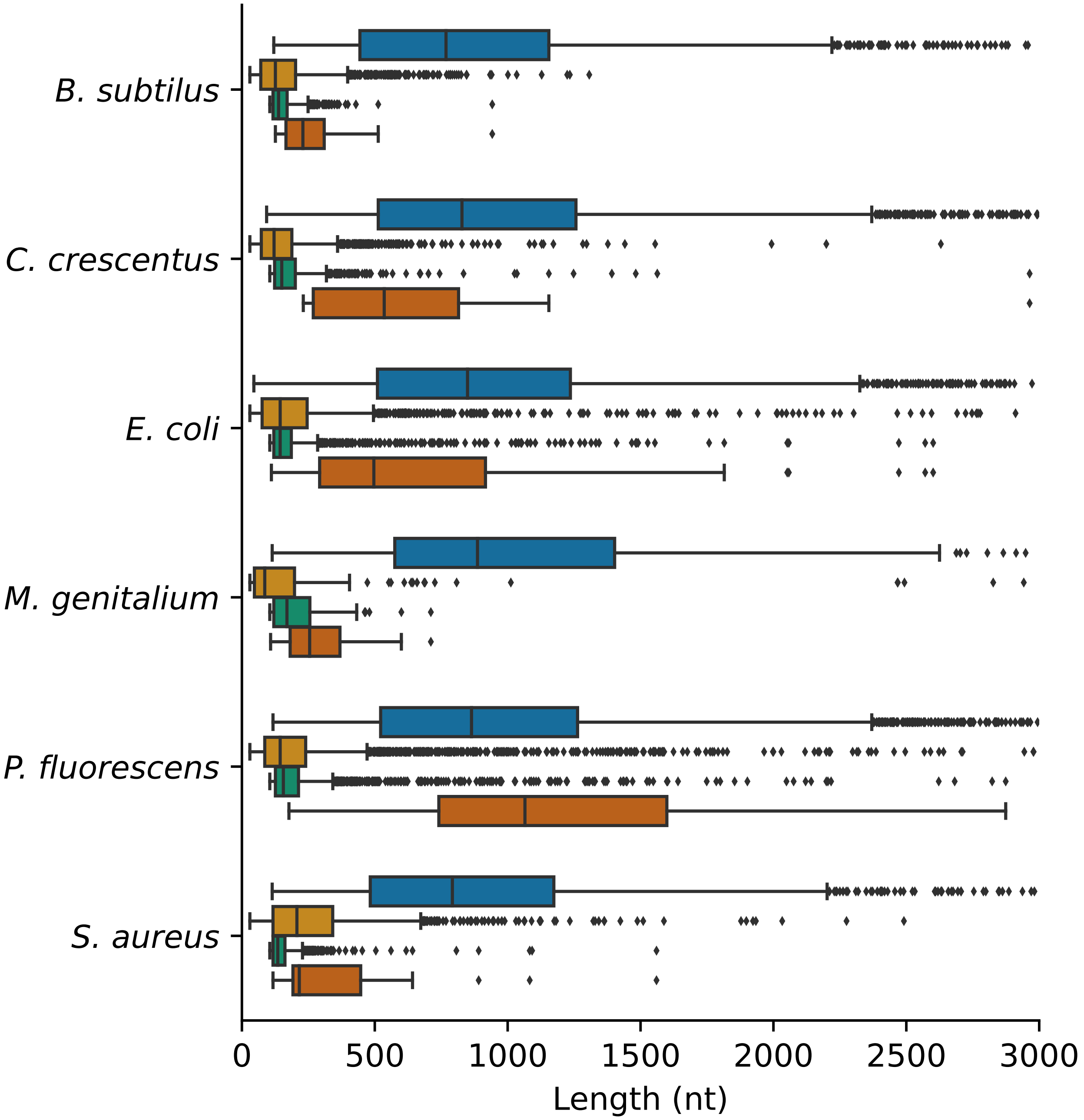


Figure 4: Reported here are the nucleotide lengths of the Ensembl genes (blue), unannotated regions (URs) extracted from the Ensembl annotations of each of the six model organisms (light orange), the StORFs (Stop-ORFs) identified from the URs (green) and the StORFs which had a high sequence similarity to known protein coding genes in Swiss-Prot (>=60% bitscore) (dark orange). X axis truncated at 3,000 nt.

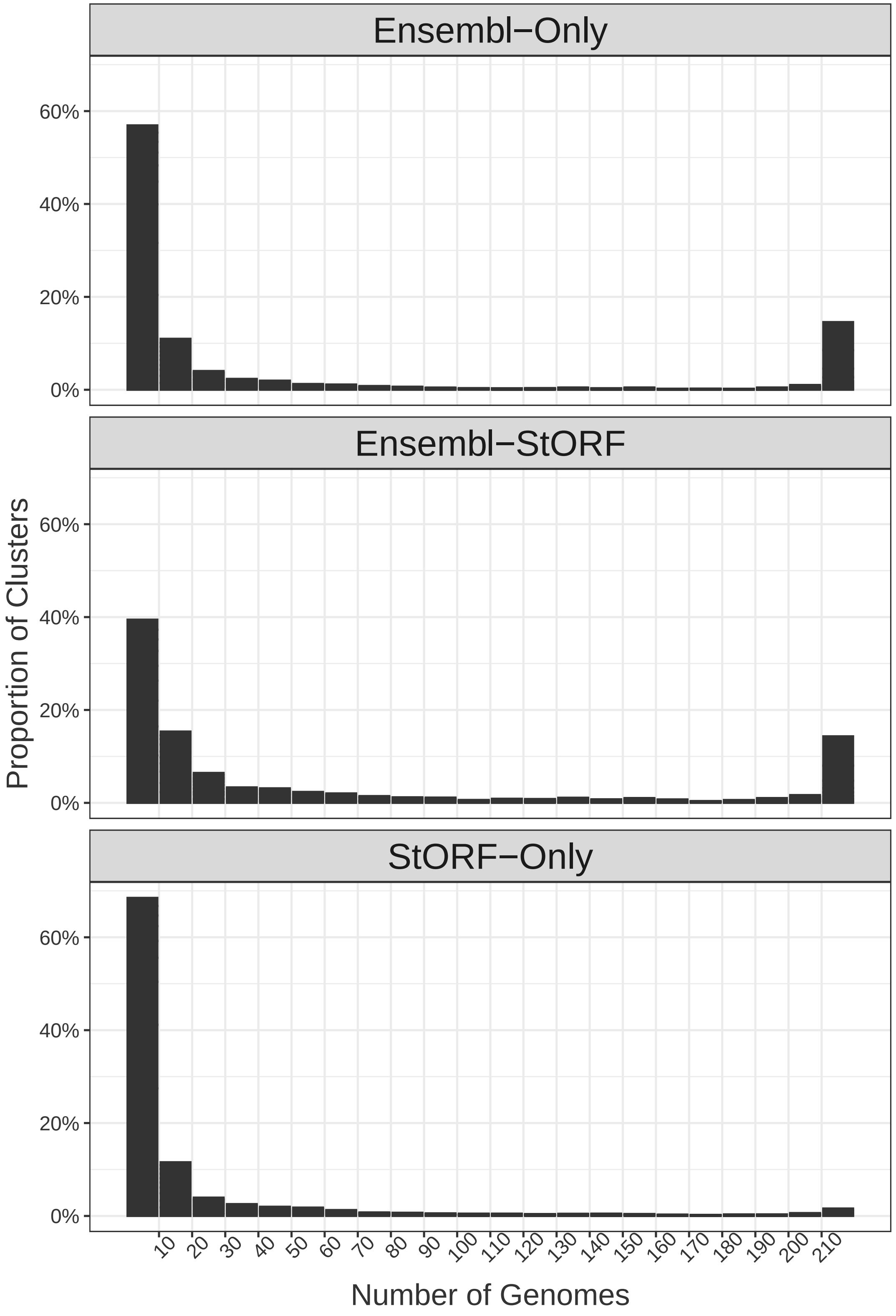


Figure 5: The distributions of gene families across the 219 *E. coli* pangenome for the Ensembl-Only, Ensembl-StORF and StORF-Only clusters. The U-shaped curve is consistent throughout the three cluster types with Ensembl-StORF containing slightly larger gene family clusters as expected due to the added StORF sequences as compared to Ensembl-Only. While the distribution is more towards the lower end for StORF-Only, a similar, albeit much less pronounced U-shaped curve is observed.

```
Reference sequence (1): Ensembl_Protein_AHM40736
Identities normalised by aligned length.
Colored by: identity

                             cov     pid   1 [        .         .         .         .         :         .         .         . 80
1 Ensembl_Protein_AHM40736 100.0% 100.0%    ------MISLPSGTRIWLVAGVTDMRKSFNGLGEQIQHVLDDNPFSGHLFIFRGRRGDTIKILWADADGLCLFTKRLEEG
2 Ensembl_Protein_AIT36070 100.0%  89.7%    ------MITLPAGSRIWLVAGITDMRNGFNGLGEQVQHVLNDNPFSGHLFIFRGRRGDTVKILWADADGLCLFTKRLEEG
3 Ensembl_Protein_AFS84250  71.6%  86.7%    ------MISLPSGTRIWLVAGVTDMRKSFNGLGEQVQHVLNDNPFSGHLFIFRGRRGDTVKILWADADGLCLFTKRLEEG
4 StORF_Sequence           100.0%  89.3%    TDREGKMISLPSGTRIWLVAGVTDMRKSFNGLGEQVQHVLNDNPFSGHLFIFRGRRGDTVKILWADADGLCLFTKRLEEG

                             cov     pid  81         .         1         .         . ] 122
1 Ensembl_Protein_AHM40736 100.0% 100.0%    QFIWPAVRDGKISITRSQLAMLLDKLDWRQPKTSRLNALTML
2 Ensembl_Protein_AIT36070 100.0%  89.7%    QFIWPAVRDGKVSITRSQLAMLLDKLDWRQPKTSRRNSLTML
3 Ensembl_Protein_AFS84250  71.6%  86.7%    LNRPGNPGD---------------------------------
4 StORF_Sequence           100.0%  89.3%    QFIWPAVRDGKVSITRSQLAMLLDKLDWRQPKTSSRNSLTML
```

Figure 6: ClustalO multiple sequence alignment of the three originally independent Ensembl representatives sequences, AHM40736, AIT36070 and AFS84250 which were clustered together by the addition of a StORF sequence which formed the single *E. coli* pangenome cluster 17,575. These three Ensembl representative sequences and their clusters have been grouped together by a single StORF sequence, thus completely changing the dynamics of this set of gene families. The sequences did not cluster together in the first round of clustering because their percentage identity was less than 90% from each other.

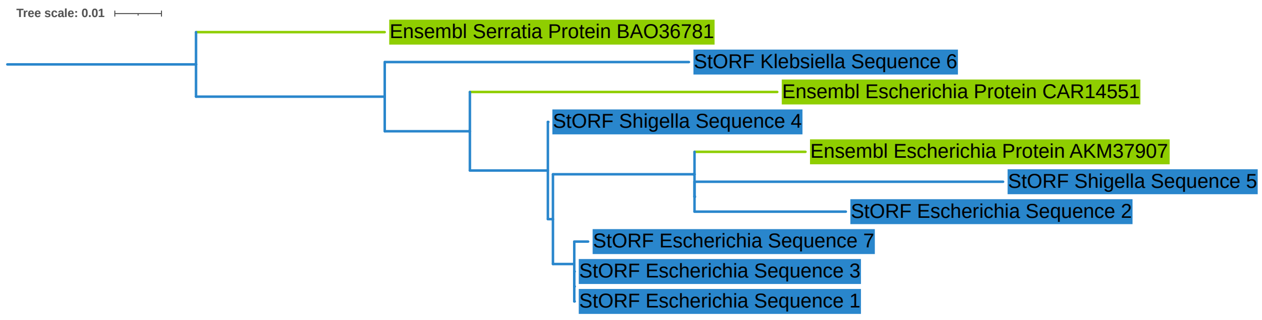


Figure 7: This is a phylogenetic tree built from the amino acid sequences of combined Cluster 124,470. This cluster consists of three Ensembl cluster representatives (clusters 1,013,244, 5364 and 382,322) and the seven StORF sequences which clustered to those representatives. This tree was created using ClustalO and Fasttree and rooted at the Ensembl_Serratia_Protein_BAO36781 sequence.

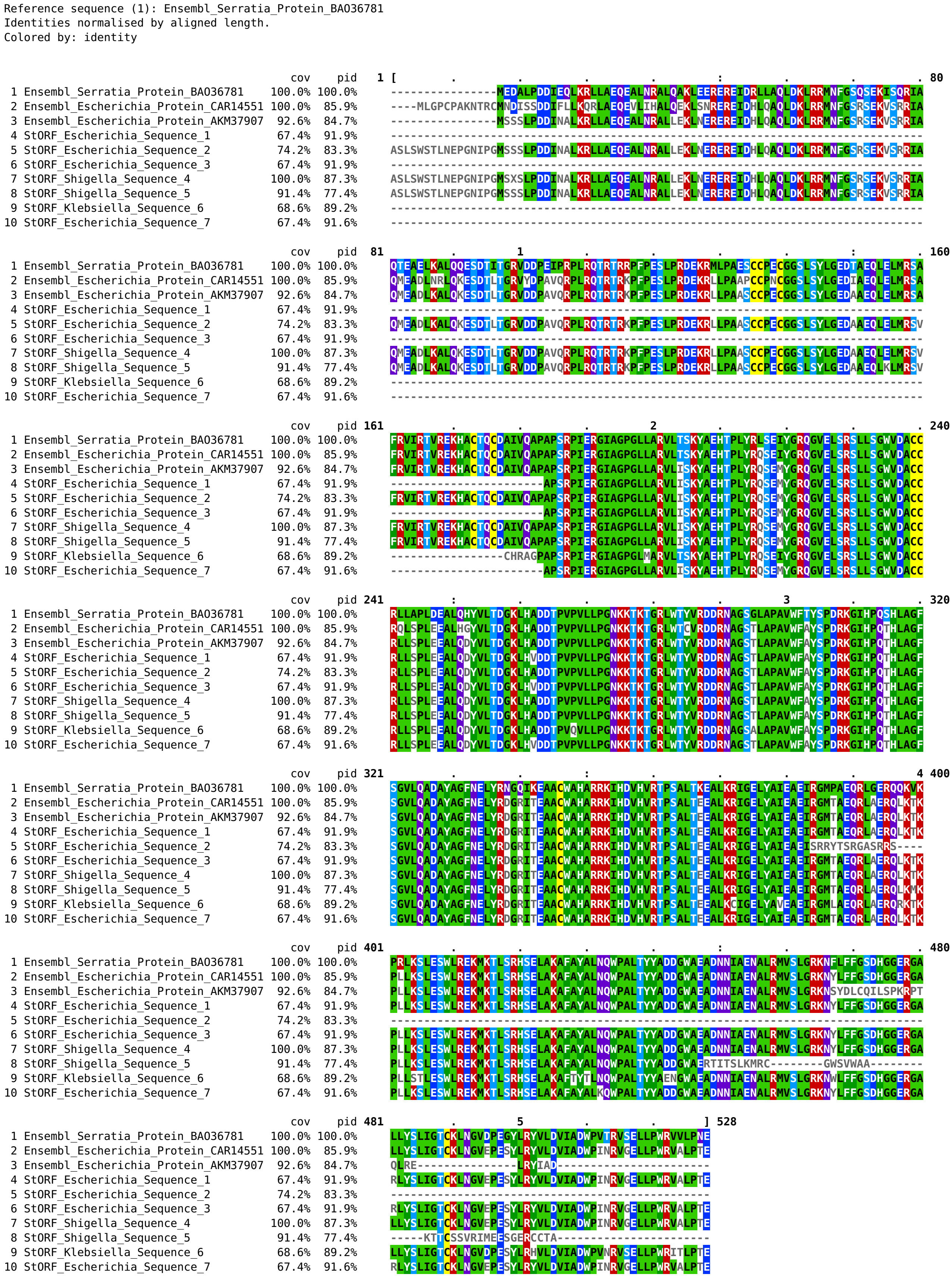


Figure 8: ClustalO multiple sequence alignment from the amino acid sequences of combined Cluster 124,470. This cluster consists of three Ensembl cluster representatives (clusters 1,013,244, 5364 and 382,322) and seven StORF sequences.

#### 2 Supplementary Tables

| | | | | | | | |
|---|---|---|---|---|---|---|---|
| Acetobacter | 14 | Clostridioides | 5 | Methanobacterium | 6 | Salmonella | 235 |
| Achromobacter | 8 | Clostridium | 79 | Methanobrevibacter | 6 | Selenomonas | 7 |
| Acidithiobacillus | 5 | Collimonas | 6 | Methanocaldococcus | 6 | Serratia | 33 |
| Acidovorax | 6 | Corynebacterium | 110 | Methanococcus | 7 | Shewanella | 26 |
| Acinetobacter | 83 | Coxiella | 10 | Methanosarcina | 26 | Shigella | 20 |
| Actinobacillus | 8 | Cronobacter | 10 | Methylobacterium | 11 | Sinorhizobium | 10 |
| Actinomyces | 12 | Cupriavidus | 6 | Microbacterium | 13 | Sphingobium | 9 |
| Aeromicrobium | 7 | Dehalococcoides | 13 | Micromonospora | 5 | Sphingomonas | 9 |
| Aeromonas | 26 | Deinococcus | 11 | Moraxella | 8 | Sphingopyxis | 9 |
| Aggregatibacter | 8 | Desulfitobacterium | 5 | Mycobacterium | 733 | Spiroplasma | 17 |
| Agrobacterium | 8 | Desulfotomaculum | 6 | Mycoplasma | 95 | Staphylococcus | 329 |
| Alcanivorax | 5 | Desulfovibrio | 19 | Myroides | 7 | Stenotrophomonas | 14 |
| Altererythrobacter | 6 | Dickeya | 6 | Neisseria | 36 | Streptococcus | 343 |
| Alteromonas | 19 | Edwardsiella | 10 | Nitrosomonas | 6 | Streptomyces | 82 |
| Amycolatopsis | 9 | Ehrlichia | 15 | Nocardia | 5 | Sulfolobus | 21 |
| Anaplasma | 19 | Enterobacter | 82 | Nostoc | 7 | Synechococcus | 25 |
| Arcobacter | 5 | Enterococcus | 83 | Oenococcus | 5 | Synechocystis | 7 |
| Arthrobacter | 12 | Erwinia | 8 | Paenibacillus | 45 | Thermoanaerobacter | 10 |
| Azospirillum | 6 | Erythrobacter | 5 | Pandoraea | 11 | Thermococcus | 18 |
| Bacillus | 283 | Escherichia | 225 | Pantoea | 14 | Thermotoga | 13 |
| Bacteroides | 17 | Eubacterium | 12 | Paraburkholderia | 9 | Thermus | 8 |
| Bartonella | 36 | Flavobacterium | 16 | Pasteurella | 12 | Thioalkalivibrio | 5 |
| Bdellovibrio | 5 | Francisella | 59 | Pectobacterium | 10 | Treponema | 40 |
| Bibersteinia | 5 | Frankia | 5 | Pediococcus | 6 | Ureaplasma | 10 |
| Bifidobacterium | 79 | Fusobacterium | 15 | Planococcus | 9 | Veillonella | 5 |
| Bordetella | 29 | Gardnerella | 6 | Porphyromonas | 7 | Vibrio | 67 |
| Borrelia | 28 | Geobacillus | 18 | Prevotella | 14 | Weissella | 5 |
| Borreliella | 6 | Geobacter | 11 | Prochlorococcus | 16 | Wolbachia | 8 |
| Brachyspira | 9 | Haemophilus | 39 | Propionibacterium | 26 | Xanthomonas | 57 |
| Bradyrhizobium | 14 | Halomonas | 5 | Proteus | 9 | Xenorhabdus | 7 |
| Brucella | 164 | Helicobacter | 108 | Providencia | 5 | Xylella | 7 |
| Buchnera | 20 | Hymenobacter | 7 | Pseudoalteromonas | 8 | Yersinia | 66 |
| Burkholderia | 183 | Janthinobacterium | 5 | Pseudomonas | 197 | Archaeoglobus | 5 |
| Caldicellulosiruptor | 8 | Klebsiella | 149 | Psychrobacter | 9 | Blattabacterium | 8 |
| Campylobacter | 74 | Lactobacillus | 108 | Pyrobaculum | 8 | Hydrogenobaculum | 5 |
| Candidate | 5 | Lactococcus | 18 | Pyrococcus | 8 | Taylorella | 5 |
| Candidatus | 176 | Legionella | 17 | Ralstonia | 17 | Zymomonas | 7 |
| Caulobacter | 5 | Leifsonia | 8 | Rhizobium | 23 | Myxococcus | 5 |
| Cedecea | 6 | Leptolyngbya | 5 | Rhodobacter | 6 | Hyphomicrobium | 5 |
| Cellulophaga | 5 | Leptospira | 16 | Rhodococcus | 20 | Aerococcus | 6 |
| Chlamydia | 147 | Leuconostoc | 12 | Rhodopseudomonas | 7 | Cyanothece | 6 |
| Chlamydophila | 7 | Listeria | 81 | Rickettsia | 57 | Lysobacter | 5 |
| Chlorobium | 7 | Mannheimia | 15 | Riemerella | 8 | Pseudonocardia | 5 |
| Chryseobacterium | 6 | Marinobacter | 11 | Ruminococcus | 9 | Clavibacter | 5 |
| Citrobacter | 16 | Mesorhizobium | 10 | Saccharomonospora | 6 | | |

Table 1: Listed here are the 6,223 genomes of 1,417 Ensembl Bacteria species grouped into their 179 genera after filtering that were used in the cross genera study.

Escherichia_coli_o157_h7_str_ec4042.ASM18177v1
Escherichia_coli_kte50.Esch_coli_KTE50_V1
Escherichia_coli_kte91.Esch_coli_KTE91_V1
Escherichia_coli_gca_001617565.ASM161756v1
Escherichia_coli_b_str_rel606.ASM1798v1
Escherichia_coli_gca_000987875.ASM98787v1
Escherichia_coli_gca_001183685.ASM118368v1
Escherichia_coli_ko11.ASM14785v3
Escherichia_coli_kte197.Esch_coli_KTE197_V1
Escherichia_coli_kte165.Esch_coli_KTE165_V1
Escherichia_coli_gca_001677475.ASM167747v1
Escherichia_coli_536.ASM1330v1
Escherichia_coli_bw25113.ASM75055v1
Escherichia_coli_k_12_gca_000974885.ASM97488v1
Escherichia_coli_53638.ASM16791v2
Escherichia_coli_str_k_12_substr_w3110.ASM1024v1
Escherichia_coli_iai39.ASM2634v1
Escherichia_coli_kte54.Esch_coli_KTE54_V1
Escherichia_coli_gca_000931565.ASM93156v1
Escherichia_coli_gca_001039415.ASM103941v1
Escherichia_coli_bl21_gold_de3_plyss_ag_.ASM2366v1
Escherichia_coli_kte233.Esch_coli_KTE233_V1
Escherichia_coli_bwh_40.Esch_coli_BWH_40_V1
Escherichia_coli_umea_3585_1.Esch_coli_UMEA_3585-1_V1
Escherichia_coli_o111_h_str_11128.ASM1076v1
Escherichia_coli_kte98.Esch_coli_KTE98_V1
Escherichia_coli_gca_000819645.ASM81964v1
Escherichia_coli_gca_000988465.ASM98846v1
Escherichia_coli_kte183.Esch_coli_KTE183_V1
Escherichia_coli_str_k_12_substr_mds42.ASM35018v1
Escherichia_coli_gca_001677495.ASM167749v1
Escherichia_coli_kte135.Esch_coli_KTE135_V1
Escherichia_coli_o157_h7_str_ss52.ASM80370v1
Escherichia_coli_str_k_12_substr_mg1655_gca_000801205.ASM80120v1
Escherichia_coli_gca_000830035.ASM83003v1
Escherichia_coli_gca_000988385.ASM98838v1
Escherichia_coli_gca_001043215.ASM104321v1
Escherichia_coli_k_12_gca_000981485.EcoliK12AG100
Escherichia_coli_uti89.ASM1326v1
Escherichia_coli_gca_000814145.ASM81414v2
Escherichia_coli_gca_001007915.ASM100791v1
Escherichia_coli_apec_o1.ASM1484v1
Escherichia_coli_gca_001030485.Esch_coli_BIDMC98_V1
Escherichia_coli_k_12_gca_000974865.ASM97486v1
Escherichia_coli_o7_k1_str_ce10.ASM22762v1
Escherichia_coli_kte118.Esch_coli_KTE118_V1
Escherichia_coli_o145_h28_str_rm13514.ASM52003v1
Escherichia_coli_bidmc_20b.Esch_coli_BIDMC_20B_V1
Escherichia_coli_o157_h7_str_tw14359.ASM2222v1
Escherichia_coli_kte229.Esch_coli_KTE229_V1
Escherichia_coli_jj1886.ASM49375v1
Escherichia_coli_kte155.Esch_coli_KTE155_V1
Escherichia_coli_dh1.ASM2336v1
Escherichia_coli_kly.ASM72530v1
Escherichia_coli_gca_000988445.ASM98844v1
Escherichia_coli_iai1.ASM2626v1
Escherichia_coli_kte29.Esch_coli_KTE29_V1
Escherichia_coli_gca_001672015.ASM167201v1
Escherichia_coli_str_st540_gca_000599625.ASM59962v1
Escherichia_coli_apec_imt5155.ASM81316v1
Escherichia_coli_o157_h7_str_ss17.ASM73034v1
Escherichia_coli_k_12_gca_000974505.ASM97450v1
Escherichia_coli_acn001.ASM105113v1
Escherichia_coli_kte67.Esch_coli_KTE67_V1
Escherichia_coli_hvh_145_4_5672112_.Esch_coli_HVH_145_4-5672112_V1
Escherichia_coli_o157_h7_str_ec4115.ASM2112v1
Escherichia_coli_k_12_gca_000974535.ASM97453v1
Escherichia_coli_gca_001030405.Esch_coli_MGH108_V1
Escherichia_coli_o139_h28_str_e24377a.ASM1774v1
Escherichia_coli_kte209.Esch_coli_KTE209_V1
Escherichia_coli_kte6.Esch_coli_KTE6_V1
Escherichia_coli_o145_h28_str_rm13516.ASM52005v1
Escherichia_coli_str_ucd_ja17_pb.UCD_JA17_pb
Escherichia_coli_chi7122.ASM30720v1
Escherichia_coli_bwh_34.Esch_coli_BWH_34_V1
Escherichia_coli_str_ucd_ja65_pb.JA65_HGAP2
Escherichia_coli_hs.ASM1776v1
Escherichia_coli_hvh_203_4_3126218_.Esch_coli_HVH_203_4-3126218_V1
Escherichia_coli_nissle_1917_gca_000714595.ASM71459v1
Escherichia_coli_o25b_h4_st131.EC958.v1
Escherichia_coli_gca_000988355.ASM98835v1
Escherichia_coli_k_12_gca_000974575.ASM97457v1
Escherichia_coli_kte79.Esch_coli_KTE79_V1
Escherichia_coli_k_12_gca_000974465.ASM97446v1
Escherichia_coli_gca_000784925.ASM78492v1
Escherichia_coli_gca_000801185.ASM80118v2
Escherichia_coli_atcc_8739.ASM1938v1
Escherichia_coli_umnk88.ASM21271v2
Escherichia_coli_ihe3034.ASM2574v1
Escherichia_coli_gca_000801165.ASM80116v1
Escherichia_coli_1303.ASM82998v1
Escherichia_coli_str_clone_d_i14_.ASM23389v1
Escherichia_coli_bl21_de3_gca_000022665.ASM2266v1
Escherichia_coli_s88.ASM2628v1
Escherichia_coli_gca_001612475.ASM161247v1
Escherichia_coli_kte45.Esch_coli_KTE45_V1
Escherichia_coli_str_sanji.ASM161075v1
Escherichia_coli_o55_h7_str_cb9615.ASM2516v1
Escherichia_coli_o91_h21_str_2009c_3740.Ec2009C-3740
Escherichia_coli_o104_h4_str_2009el_2071.ASM29947v1
Escherichia_coli_gca_000833145.ASM83314v1
Escherichia_coli_gca_000952955.EcRV308Chr
Escherichia_coli_atcc_25922_gca_000743255.ASM74325v1
Escherichia_coli_bl21_de3_.ASM956v1
Escherichia_coli_kte65.Esch_coli_KTE65_V1
Escherichia_coli_kte212.Esch_coli_KTE212_V1
Escherichia_coli_o26_h11_str_11368.ASM9100v1
Escherichia_coli_kte173.Esch_coli_KTE173_V1
Escherichia_coli_gca_001721525.ASM172152v1
Escherichia_coli_gca_001420955.ASM142095v1
Escherichia_coli_str_st540_gca_000599645.ASM59964v1
Escherichia_coli_bw2952.ASM2234v1
Escherichia_coli_rs218.ASM80084v2
Escherichia_coli_na114.ASM21476v2
Escherichia_coli_abu_83972.ASM14836v1
Escherichia_coli_gca_000971615.ASM97161v1
Escherichia_coli_kte192.Esch_coli_KTE192_V1
Escherichia_coli_hvh_193_4_3331423_.Esch_coli_HVH_193_4-3331423_V1
Escherichia_coli_k_12_gca_000974825.ASM97482v1
Escherichia_coli_gca_001183665.ASM118366v1
Escherichia_coli_umnf18.ASM22000v2
Escherichia_coli_kte158.Esch_coli_KTE158_V1
Escherichia_coli_kte84.Esch_coli_KTE84_V1
Escherichia_coli_hvh_199_4_5670322_.Esch_coli_HVH_199_4-5670322_V1
Escherichia_coli_str_clone_d_i2_.ASM23387v1
Escherichia_coli_ecc_1470_gca_000831565.ASM83156v1
Escherichia_coli_kte19.Esch_coli_KTE19_V1
Escherichia_coli_kte42.Esch_coli_KTE42_V1
Escherichia_coli_kte161.Esch_coli_KTE161_V1
Escherichia_coli_kte46.Esch_coli_KTE46_V1
Escherichia_coli_kte227.Esch_coli_KTE227_V1
Escherichia_coli_gca_001721125.ASM172112v1
Escherichia_coli_bwh_24.Esch_coli_BWH_24_V1_PacBio
Escherichia_coli_pcn061.ASM102912v1
Escherichia_coli_str_ucd_ja23_pb.UCD_JA23_pb
Escherichia_coli_o157_h7_str_edl933.ASM666v1
Escherichia_coli_gca_000953515.EcHMS174Chr
Escherichia_coli_str_k_12_substr_dh10b.ASM1942v1
Escherichia_coli_str_st540_gca_000599665.ASM59966v1
Escherichia_coli_kte218.Esch_coli_KTE218_V1
Escherichia_coli_kte222.Esch_coli_KTE222_V1
Escherichia_coli_hvh_147_4_5893887_.Esch_coli_HVH_147_4-5893887_V1
Escherichia_coli_cft073.ASM744v1
Escherichia_coli_lf82.ASM28449v1
Escherichia_coli_pmv_1.EcoPMV1
Escherichia_coli_hvh_195_3_7155360_.Esch_coli_HVH_195_3-7155360_V1
Escherichia_coli_gca_000988425.ASM98842v1
Escherichia_coli_ko11fl.ASM25802v1
Escherichia_coli_o157_h7_str_edl933_gca_000732965.ASM73296v1
Escherichia_coli_vr50.ASM96851v1
Escherichia_coli_p12b.ASM25727v1
Escherichia_coli_xuzhou21.ASM26212v1
Escherichia_coli_kte154.Esch_coli_KTE154_V1
Escherichia_coli_gca_001518955.ASM151895v1
Escherichia_coli_kte123.Esch_coli_KTE123_V1
Escherichia_coli_kte235.Esch_coli_KTE235_V1
Escherichia_coli_hvh_140_4_5894387_.Esch_coli_HVH_140_4-5894387_V1
Escherichia_coli_umea_3662_1.Esch_coli_UMEA_3662-1_V1
Escherichia_coli_b41.ASM19470v2
Escherichia_coli_55989.ASM2624v1
Escherichia_coli_hvh_141_4_5995973_.Esch_coli_HVH_141_4-5995973_V1
Escherichia_coli_kte213.Esch_coli_KTE213_V1
Escherichia_coli_gca_001515725.ASM151572v1
Escherichia_coli_gca_001750845.ASM175084v1
Escherichia_coli_k_12_gca_000974405.ASM97440v1
Escherichia_coli_dh1_gca_000270105.ASM27010v1
Escherichia_coli_sms_3_5.ASM1964v1
Escherichia_coli_kte9.Esch_coli_KTE9_V1
Escherichia_coli_etec_h10407.ASM21047v1
Escherichia_coli_o145_h28_str_rm12761.ASM66239v1
Escherichia_coli_o103_h2_str_12009.ASM1074v1
Escherichia_coli_apec_o78.ASM33275v1
Escherichia_coli_aa86.ASM21139v1
Escherichia_coli_kte147.Esch_coli_KTE147_V1
Escherichia_coli_bidmc_19c.Esch_coli_BIDMC_19C_V1_PacBio
Escherichia_coli_w_gca_000258145.ASM25814v1
Escherichia_coli.HUSEC2011CHR1
Escherichia_coli_gca_001183645.ASM118364v1
Escherichia_coli_o157_h7_str_tw14588.ASM15512v1
Escherichia_coli_o83_h1_str_nrg_857c.ASM18334v1
Escherichia_coli_fap1.ASM76543v1
Escherichia_coli_k_12.ASM80076v1
Escherichia_coli_gca_001030345.Esch_coli_MGH107_V1
Escherichia_coli_kte184.Esch_coli_KTE184_V1
Escherichia_coli_o157_h7_str_sakai.ASM886v1
Escherichia_coli_str_k_12_substr_mg1655.ASM584v2
Escherichia_coli_umn026.ASM2632v2
Escherichia_coli_str_k_12_substr_mc4100.MYMC4100
Escherichia_coli_str_st540_gca_000599705.ASM59970v1
Escherichia_coli_o55_h7_str_rm12579.ASM24551v1
Escherichia_coli_kte17.Esch_coli_KTE17_V1
Escherichia_coli_gca_001280345.ASM128034v1
Escherichia_coli_er2796.ASM80021v1
Escherichia_coli_96_154.ASM21524v2
Escherichia_coli_se11.ASM1038v1
Escherichia_coli_kte140.Esch_coli_KTE140_V1
Escherichia_coli_kte115.Esch_coli_KTE115_V1
Escherichia_coli_um146.ASM14860v1
Escherichia_coli_str_st540.ASM59784v1
Escherichia_coli_b7a_gca_000725265.ASM72526v1
Escherichia_coli_w_gca_000184185.ASM18418v1
Escherichia_coli_kte203.Esch_coli_KTE203_V1
Escherichia_coli_gca_001675145.ASM167514v1
Escherichia_coli_ly180.ASM46851v1
Escherichia_coli_o145_h28_str_rm12581.ASM67129v1
Escherichia_coli_pcn033.ASM21951v3
Escherichia_coli_ed1a.ASM2630v1
Escherichia_coli_kte196.Esch_coli_KTE196_V1
Escherichia_coli_o104_h4_str_2011c_3493.ASM29945v1
Escherichia_coli_kte100.Esch_coli_KTE100_V1
Escherichia_coli_str_st540_gca_000599685.ASM59968v1
Escherichia_coli_kte56.Esch_coli_KTE56_V1
Escherichia_coli_o104_h4_str_2009el_2050.ASM29925v1
Escherichia_coli_hvh_90_4_3191362_.Esch_coli_HVH_90_4-3191362_V1
Escherichia_coli_o127_h6_str_e2348_69.ASM2654v1
Escherichia_coli_kte43.Esch_coli_KTE43_V1
Escherichia_coli_gca_001645235.ASM164523v1
Escherichia_coli_kte190.Esch_coli_KTE190_V1
Escherichia_coli_se15.ASM1048v1

Table 2: The names of each of the 219 *Escherichia coli* genomes used in the pangenome analysis.

| Model Organism | # Genes | # URs | Longest UR | Mean / Median UR Length [**SD**] |
|---|---|---|---|---|
| *B. subtilis* | 4,133 | 2,711 | 1,307 | 161.80/126.00 [**137.73**] |
| *C. crescentus* | 3,875 | 2,321 | 3,377 | 160.41/121.00 [**172.78**] |
| *E. coli* | 4,257 | 2,743 | 6,175 | 225.73/144.00 [**353.32**] |
| *M. genitalium* | 559 | 157 | 4,822 | 290.04/86.00 [**673.23**] |
| *P. fluorescens* | 5,266 | 3,509 | 19,988 | 261.81/144.00 [**633.71**] |
| *S. aureus* | 2,556 | 1,666 | 2,491 | 262.87/207.00 [**235.19**] |

Table 3: The results of running UR-Extractor on the Ensembl annotations for the six model organisms. Lengths presented are in nt and are without the 50nt extension at each end. Standard deviation is abbreviated as [**SD**].

| Model Organism | # Genes | # URs | Longest UR | Mean / Median UR Length [**SD**] |
|---|---|---|---|---|
| *B. subtilis* | 4,016 | 2,619 | 6,159 | 182.78/125.00 [**311.45**] |
| *C. crescentus* | 3,704 | 2,394 | 6,494 | 182.41/131.00 [**250.03**] |
| *E. coli* | 4,263 | 2,743 | 3,955 | 205.00/142.00 [**247.20**] |
| *M. genitalium* | 995 | 636 | 2,546 | 205.92/133.00 [**221.21**] |
| *P. fluorescens* | 5,421 | 3,524 | 4,164 | 193.42/139.00 [**239.26**] |
| *S. aureus* | 2,534 | 1,650 | 12,232 | 274.03/203.00 [**492.37**] |

Table 4: This table presents the results of running UR-Extractor on the Prodigal CDS predictions for the six model organisms. Lengths presented are in nt and are without the 50nt extension at each end. Standard deviation is abbreviated as [**SD**].

| Model Organism | # StORFs | Recovered [**Non-vitiated**] |
|---|---|---|
| *B. subtilis* | 2,472 | 12 [**51**] |
| *C. crescentus* | 1,827 | 26 [**100**] |
| *E. coli* | 2,866 | 22 [**72**] |
| *M. genitalium* | 587 | 1 [**6**] |
| *P. fluorescens* | 3,227 | 14 [**51**] |
| *S. aureus* | 2,123 | 6 [**16**] |

Table 5: This table contains the number of Prodigal StORFs and the number of non-vitiated Ensembl genes recovered by StORF-Reporter which Prodigal missed. Non-vitiated genes are those which had an overlap of less than 50 nt with a Prodigal predicted CDS, thus allowing for them to be included in an extracted UR.

| Model Organism | Swiss-Prot Hits [**Subject Hit >=80%**] | Intra-Genome Hits [**Subject Hit >=80%**] |
|---|---|---|
| *B. subtilis* | 45 [**30**] | 39 [**33**] |
| *C. crescentus* | 6 [**4**] | 60 [**48**] |
| *E. coli* | 75 [**52**] | 34 [**29**] |
| *M. genitalium* | 184 [**5**] | 182 [**2**] |
| *P. fluorescens* | 16 [**5**] | 42 [**35**] |
| *S. aureus* | 19 [**1**] | 25 [**13**] |

Table 6: The table contains the number of Prodigal StORFs which were reported with a hit to either the SwissProt or Intra-Genome protein database. Intra-Genome is the proteome of the same model organism. DIAMOND blastp hits are recorded with a minimum of a 60 bit score and in bold are reported with a subject coverage of 80%.

| Data | Unannotated Regions | StORFs |
|---|---|---|
| Number of Sequences | 673,136 | 652,056 |
| Median Number Per Genome | 3,038 | 2,958 |
| Longest Sequence (nt) | 45,683 | 14,334 |
| Median Sequence Length (nt) **[Std]** | 234 **[292.97]** | 141 **[177.48]** |

Table 7: The numbers and lengths of the unannotated regions (URs) and StORFs extracted from the 219 *Escherichia coli* genomes are presented here. Although there was variability in the genome quality across this set of genomes, the numbers reported here are similar to those reported for the 6 model organisms.

| Data | Unannotated Regions | StORFs |
|---|---|---|
| Number of Sequences | 14,221,482 | 13,301,175 |
| Median Number Per Genome | 2,305 | 1,981 |
| Longest Sequence (nt) | 86,235 | 47,790 |
| Median Sequence Length (nt) **[Std]** | 240 **[366.07]** | 147 **[213.24]** |

Table 8: The numbers and lengths of unannotated regions (UR) and StORFs extracted from the 6,223 genomes of Ensembl Bacteria are presented here. While there was variability in the genome quality across this set of genomes, the numbers reported here are similar to those reported for the 6 model organisms and the *E. coli* pangenome analysis.

| | Genome Triplet Abundance | | | Prodigal Gene Stop Usage | | | | Prodigal StORF Stop Usage | | | |
|---|---|---|---|---|---|---|---|---|---|---|---|
| Genomes | TGA [%] | TAG [%] | TAA [%] | TGA [%] | TAG [%] | TAA [%] | $\tilde{\chi}^2$ p-value | TGA [%] | TAG [%] | TAA [%] | $\tilde{\chi}^2$ p-value |
| *B. subtilis* | 180,347 **[49.57]** | 52,378 **[14.39]** | 131,084 **[36.03]** | 932 **[23.21]** | 563 **[14.02]** | 2,521 **[62.77]** | <0.00001 | 1,025 **[41.46]** | 424 **[17.15]** | 1,023 **[41.38]** | <0.00001 |
| *C. crescentus* | 102,367 **[69.85]** | 32,635 **[22.27]** | 11,541 **[7.87]** | 1,735 **[46.84]** | 1,230 **[33.21]** | 739 **[19.95]** | <0.00001 | 1,059 **[57.96]** | 458 **[25.07]** | 310 **[16.97]** | <0.00001 |
| *E. coli* | 164,560 **[46.64]** | 53,119 **[15.05]** | 135,187 **[38.31]** | 1,232 **[28.90]** | 332 **[7.79]** | 2,699 **[63.31]** | <0.00001 | 1,093 **[38.14]** | 486 **[16.96]** | 1,287 **[44.91]** | <0.00001 |
| *M. genitalium* | 25,382 **[28.26]** | 18,982 **[21.13]** | 45,456 **[50.60]** | 612 **[61.51]** | 110 **[11.06]** | 273 **[27.44]** | <0.00001 | 261 **[44.46]** | 100 **[17.04]** | 226 **[38.50]** | <0.00001 |
| *P. fluorescens* | 189,251 **[64.36]** | 56,411 **[19.18]** | 48,377 **[16.45]** | 3,010 **[55.52]** | 770 **[14.20]** | 1,641 **[30.27]** | <0.00001 | 1,644 **[50.95]** | 750 **[23.24]** | 833 **[25.81]** | <0.00001 |
| *S. aureus* | 119,798 **[30.87]** | 73,821 **[19.02]** | 194,474 **[50.11]** | 268 **[10.58]** | 379 **[14.96]** | 1,886 **[74.46]** | <0.00001 | 512 **[24.12]** | 429 **[20.21]** | 1,182 **[55.68]** | <0.00001 |

Table 9: Presented in this table are the following: the triplet abundance of the three canonical stop codons found throughout the six model organism genomes (totaled from both forward and reverse strands), the stop codons used in the Prodigal predicted CDS genes, and the end stop codon used in the StORFs identified from within the URs reported by Prodigal, both from the 6 model organisms which have been inspected. A chi squared test was performed on each model organism: triplet abundance vs Prodigal gene stop codon usage and triplet abundance vs StORF stop codon. Each test resulted in a rounded p-value of <0.00001.

| Genomes | Genome Triplet Abundance | | | Ensembl Gene Stop Usage | | | | Ensembl StORF Stop Usage | | | |
|---|---|---|---|---|---|---|---|---|---|---|---|
| | TGA [%] | TAG [%] | TAA [%] | TGA [%] | TAG [%] | TAA [%] | $\tilde{\chi}^2$ p-value | TGA [%] | TAG [%] | TAA [%] | $\tilde{\chi}^2$ p-value |
| *B. subtilis* | 180,347 **[49.57]** | 52,378 **[14.39]** | 131,084 **[36.03]** | 927 **[23.11]** | 560 **[13.96]** | 2,524 **[62.93]** | <0.00001 | 951 **[40.96]** | 374 **[16.11]** | 997 **[42.94]** | <0.00001 |
| *C. crescentus* | 102,367 **[69.85]** | 32,635 **[22.27]** | 11,541 **[7.87]** | 1,770 **[47.36]** | 1,225 **[32.78]** | 742 **[19.86]** | <0.00001 | 916 **[58.94]** | 391 **[25.16]** | 247 **[15.89]** | <0.00001 |
| *E. coli* | 164,560 **[46.64]** | 53,119 **[15.05]** | 135,187 **[38.31]** | 1,151 **[28.41]** | 279 **[6.89]** | 2,621 **[64.70]** | <0.00001 | 1,096 **[39.17]** | 437 **[15.62]** | 1,265 **[45.21]** | <0.00001 |
| *M. genitalium* | 25.382 **[28.26]** | 18.982 **[21.13]** | 45.456 **[50.60]** | 0 **[0]** | 129 **[27.10]** | 347 **[72.90]** | <0.00001 | 80 **[47.62]** | 20 **[11.90]** | 68 **[40.48]** | <0.00001 |
| *P. fluorescens* | 189,251 **[64.36]** | 56,411 **[19.18]** | 48,377 **[16.45]** | 2,869 **[55.41]** | 734 **[14.18]** | 1,575 **[30.42]** | <0.00001 | 1,756 **[51.58]** | 761 **[22.35]** | 887 **[26.06]** | <0.00001 |
| *S. aureus* | 119,798 **[30.87]** | 73,821 **[19.02]** | 194,474 **[50.11]** | 271 **[10.84]** | 379 **[15.16]** | 1,850 **[74.00]** | <0.00001 | 477 **[23.23]** | 370 **[18.02]** | 1,206 **[58.74]** | <0.00001 |

Table 10: Presented in this table are the following: the triplet abundance of the three canonical stop codons found throughout the six model organism genomes (totaled from both forward and reverse strands), the stop codons used in the Ensembl annotated CDS genes, and both end stop codons used in the StORFs identified from within the URs reported by Ensembl, both from the 6 model organisms which have been inspected. A chi squared test was performed on each model organism: triplet abundance vs Ensembl gene stop codon usage and triplet abundance vs StORF stop codon. Each test resulted in a rounded p-value of <0.00001.

### 3 Processing

Listed below are the parameters used in the extraction of URs, StORFs and CD-Hit sequence clustering. The full set of parameters for UR_Extractor and StORF_Finder, including the default options which were not modified are available on the StORF-Reporter Github repository (`https://github.com/NickJD/StORF-Reporter`)

#### 3.1 UR_Extractor User Menu

```
python3 UR_Extractor.py -f Ensembl_Genome.fasta -gff Ensembl_Genome.gff3  -o
    Ensembl_Genome_UR -gz True
```

Listing 1: Example parameter set used for UR_Extractor from all Ensembl genomes.

#### 3.2 StORF_Finder User Menu

```
python3 StORF_Finder.py -f Ensembl_Genome_UR.fasta -aa True -gff True -o
    Ensembl_Genome_UR_StORFs
```

Listing 2: Example parameter set used for UR_Extractor from all Ensembl genomes.

#### 3.3 CD-Hit Clustering

```
cd-hit -i  Escherichia_coli_PEP.fa -o Escherichia_coli_PEP.fa_CD_c90_s60 -s 0.6 -c 0.9
-sc 1 -sf 1 -p 1 -g 1 -d 0 -M 10000 -T 8
```

Listing 3: Listed here are the parameters used for the CD-Hit sequence clustering. Each clustering round was performed with the same parameters for both the Ensembl representatives and the inclusion of the StORF sequences, for both the cross-genera and *E. coli* pangenome analysis.
